# Investigating the shared genetic architecture between multiple sclerosis and inflammatory bowel diseases

**DOI:** 10.1101/2020.11.16.385914

**Authors:** Yuanhao Yang, Hannah Musco, Steve Simpson-Yap, Zhihong Zhu, Ying Wang, Xin Lin, Jiawei Zhang, Bruce Taylor, Jacob Gratten, Yuan Zhou

**Affiliations:** Mater Research, Translational Research Institute, Brisbane, QLD, Australia; Institute for Molecular Bioscience, The University of Queensland, Brisbane, QLD, Australia; Menzies Institute for Medical Research, University of Tasmania, Hobart, TAS, Australia; Neuroepidemiology Unit, Melbourne School of Population & Global Health, The University of Melbourne, Melbourne, VIC, Australia; National Centre for Register-based Research, Aarhus University, Aarhus, Denmark; Department of General Surgery and Department of Gastrointestinal Surgery, the First Affiliated Hospital of Anhui Medical University, Hefei, China

## Abstract

An epidemiological association between multiple sclerosis (MS) and inflammatory bowel disease (IBD) is well-established, but whether this reflects a shared genetic aetiology, and whether consistent genetic relationships exist between MS and the two predominant subtypes of IBD, ulcerative colitis (UC) and Crohn’s disease (CD), remains unclear. Here, we used genome-wide association study (GWAS) summary data to estimate genetic correlations *(r_g_)* between MS and each of IBD, UC and CD, finding that the *r_g_* between MS and UC was approximately twice that between MS and CD. On the basis of these genetic correlations, we performed cross-trait meta-analysis of GWAS summary data for MS and each of IBD, UC and CD, identifying a total of 42 novel SNPs shared between MS and IBD (N=19), UC (N=14), and CD (N=18). We then used multiple Mendelian randomization (MR) methods to investigate causal relationships between these diseases, finding suggestive but inconclusive evidence for a causal effect of MS on UC and IBD, and no or weak and inconsistent evidence for a causal effect of IBD or UC on MS. There was also no evidence for causality in bidirectional analyses of MS and CD. We also investigated tissue- and cell-type-specific enrichment of SNP heritability for each disease using stratified LD score regression. At the tissue level, we observed largely consistent patterns of enrichment for all four diseases in immune system-related tissues, including lung, spleen and whole blood, and in contrast to prior studies, small intestine. At the cell-type level, we identified significant enrichment for all diseases in CD4^+^ T cells in lung, and for MS, IBD and CD in CD8^+^ cytotoxic T cells in both lung and spleen, and regulatory T cells in lung. Our study sheds new insights into the biological basis of comorbidity between MS and both UC and CD.

## Introduction

Multiple sclerosis (MS) is a complex autoimmune disease of the central nervous system (CNS) involving demyelination of neurons and subsequent neurodegeneration^1^. Inflammatory bowel disease (IBD) is characterized by chronic inflammation of the gastrointestinal (GI) tract, and encompasses both ulcerative colitis (UC; inflammation predominantly in the large intestine and rectum, occasionally in the terminal ileum) and Crohn’s disease (CD; inflammation in any part of the GI tract)^2^. Evidence for reciprocal comorbidity of MS and IBD has grown in recent years^3–5^. For example, a large meta-analysis^6^ with over one million participants from MS and IBD registries found that MS was associated with a 55% increased risk of IBD, and reciprocally, that IBD patients had a 53% increased risk of MS. No differences in MS prevalence between patients with UC or CD were detected in that study, but others^7,8^ have reported greater risk of MS in UC patients, and vice versa, compared to those with CD.

Both MS and IBD are moderately heritable, with estimated liability-scale single nucleotide polymorphism (SNP) heritability of 19%^9^ and around 25% (27% for UC and 21% for CD)^10^, respectively. Large-scale case-control genome-wide association studies (GWAS) for MS, IBD (case samples including UC and CD), UC and CD have identified hundreds of variants conferring risk for each disease^9,11,12^, including some shared risk loci (e.g. *IL7R*^13,14^ and *IL2RA^13,15^).* These findings suggest that MS may have partially shared genetic risk with UC and CD, but the magnitude of the genetic overlap remains unclear, as does the question of whether any genetic overlap reflects pleiotropy or causality. Interestingly, previous studies^16,17^ have reported evidence that MS may share different genetic factors with each of UC and CD. For example, MS is genetically more similar to UC than CD in relation to the major histocompatibility complex (MHC) region^18^. Prior studies have also revealed multiple tissues (e.g. lung, spleen, peripheral blood) enriched for SNP heritability of MS, UC and CD (e.g. Finucane et al. 2018^19^, IMSGC et al 2019^9^), although further investigation is needed to determine if this shared enrichment reflects involvement of the same versus distinct cell types across diseases. Addressing these questions could help to gain a deeper understanding of the biological mechanisms underlying comorbid MS and IBD.

A dilemma that doctors face in immunology and gastroenterology clinics is how to treat patients with both MS and IBD. For example, it has been reported that the cytokine, interferon-β, used to treat MS can increase the severity of IBD symptoms^20^, and conversely, that a TNF-α antagonist agent that is effective for IBD can worsen the clinical course of MS^21^. For these reasons, an improved understanding of genetic relationships between MS and comorbid IBD may lead to safer and more effective interventions for both diseases.

In this study, we used large-scale GWAS summary data to examine genetic correlations and potential causality between MS and each of IBD, UC and CD. We performed cross-trait GWAS meta-analyses between MS and IBD, UC and CD, and identified novel genetic risk variants not previously associated with the individual traits. We integrated GWAS summary data with tissue and cell-type-specific gene expression data to determine if SNP heritability for MS and each of IBD, UC and CD is enriched in the same as opposed to distinct tissues and cell types, and we used Summary-datebased Mendelian randomisation (SMR)^22^ to identify putative functional genes shared between diseases. A flowchart of our analysis strategy is provided in Figure S1.

## Results

### Genetic correlations between MS and IBDs

We first applied linkage disequilibrium (LD) score regression (LDSC)^23^ to estimate the liability-scale SNP heritability for MS and each of IBD, UC and CD. Consistent with the literature^9,10^, the liability-scale SNP heritability (without constrained intercept) was 13% for MS, 16% for IBD, 15% for UC, and 25% for CD (Table S1). We then used bivariate LDSC to estimate genetic correlations between MS and each of IBD, UC and CD. The genetic correlation (without constrained intercept) between MS and UC (*r_g_*=0.33, p=1.66×10^−13^) was roughly twice that between MS and CD (*r_g_*=0.16, p=2.40×10^−3^), with the MS-IBD estimate intermediate between these values (*r_g_*=0.28, p=2.01×10^−10^), as expected, reflecting that the IBD GWAS case sample is comprised of both UC and CD patients (Figure 1). The intercept of genetic covariance between MS and IBD (or UC or CD) was estimated at around 0.10, indicating mild sample overlap between MS and IBD (or UC or CD). For comparison, the genetic correlation between UC and CD was 0.70 (p=2.05×10^−47^). These estimates were slightly weaker after constraining the LDSC intercept, but nonetheless all remained Bonferroni significant (p<1.25×10^−2^).

**Figure 1.**
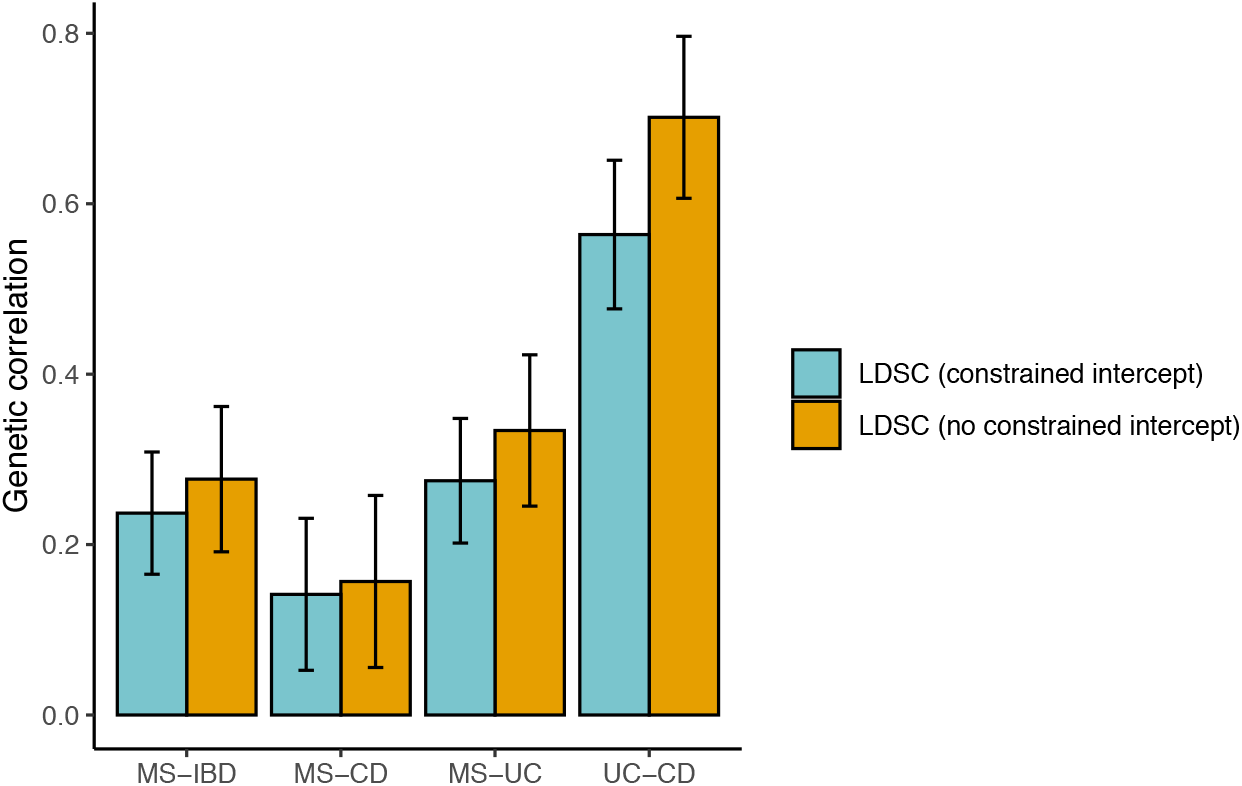
Summary of pairwise genetic correlations estimated using LD score regression with and without constrained intercept. Error bars represents the 95% confidence intervals (CIs) of the genetic correlations.

### Local genetic correlations between MS and IBDs

We used the ρ-HESS (Heritability Estimation from Summary Statistics) method^24^ to evaluate local genetic correlations across the genome between MS and each of IBD, UC and CD. In each of the three pairwise comparisons (MS-IBD, MS-UC, MS-CD), there was no evidence for a difference in local genetic correlation in regions harbouring MS-specific loci versus IBD-, UC- and CD-specific loci (Figure 2). Additionally, local genetic correlations in disease-specific loci (e.g. MS-specific and IBD-specific loci for the MS-IBD comparison) were all largely consistent with the genome-wide *r_g_*estimates from bivariate LDSC. Significant local genetic correlations were identified in the five MHC regions on chromosome 6 for MS-UC and MS-IBD, but not MS-CD, with the caveat that some of the latter estimates may be unreliable due to non-significant local SNP heritability estimates (Table S2; Figures S2-4).

**Figure 2.**
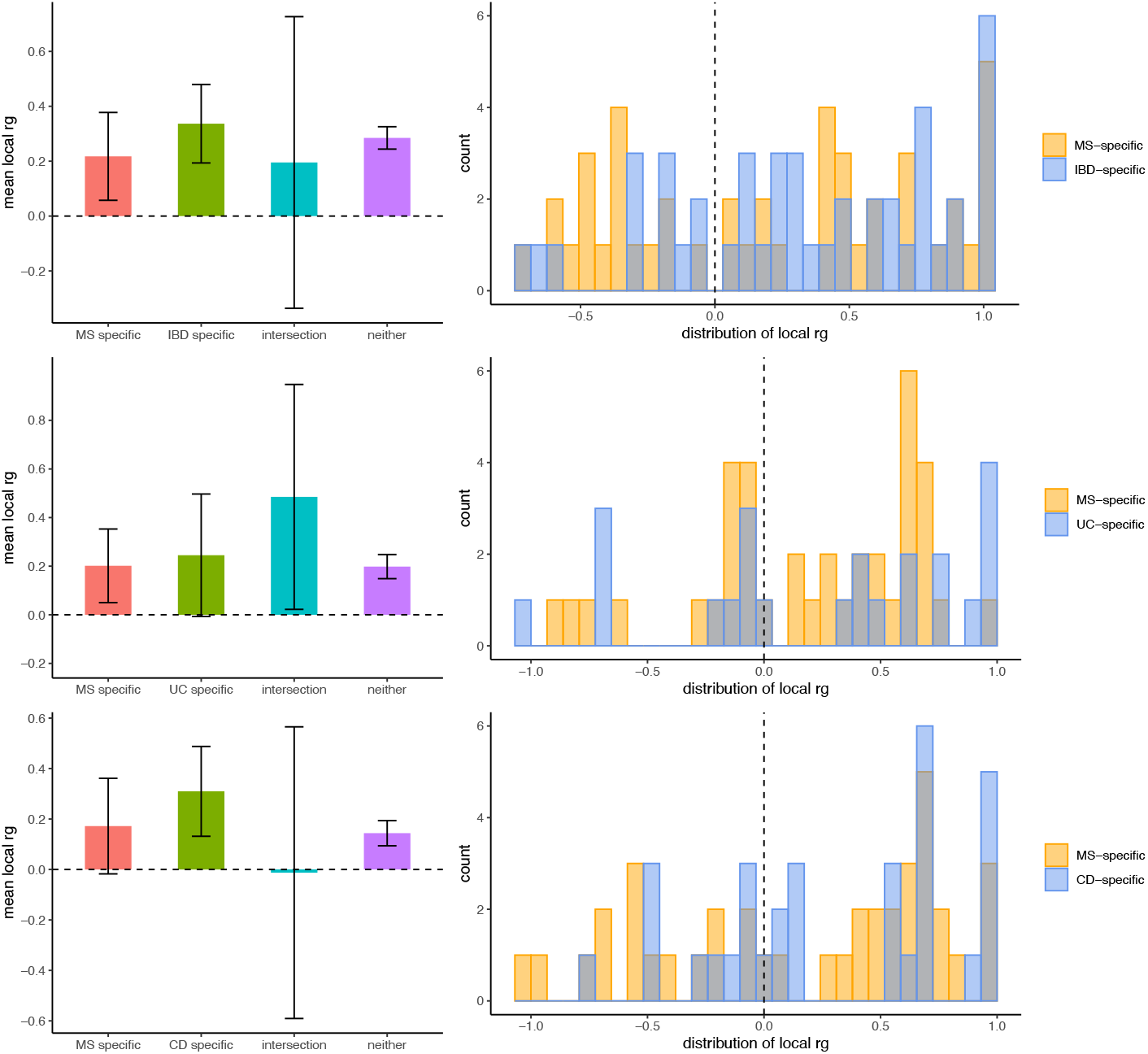
Local genetic correlations between MS and IBD, UC and CD, respectively. For each pair of diseases, local genetic correlation estimates are provided for regions harbouring disease-specific risk variants (p<5×10^−8^), regions harbouring shared risk variants (“intersection”) and all other regions (“neither”). Local genetic correlations with estimates less than −1 or greater than 1 were forced to −1 or 1, respectively. Error bars represent the 95% CIs, calculated using a jack-knife method. For MS-IBD, 40, 45, and 8 regions were included in the ‘MS-specific’, ‘IBD-specific’, and ‘intersection’ categories; for MS-UC, 38, 23, and 7 regions were included in the ‘MS-specific’, ‘UC-specific’, and ‘intersection’ categories; for MS-CD, 39, 32, and 7 regions were included in the ‘MS-specific’, ‘CD-specific’, and ‘intersection’ categories.

### Novel genetic loci from cross-trait meta-analysis of MS and each of IBD, UC, and CD

Based on evidence for significant genetic correlations between MS and each of IBD, UC and CD, we performed cross-trait meta-analyses using MTAG (Multi-Trait Analysis of GWAS)^25^. We identified 19 novel SNP loci (p<5×10^−8^; summarised in Table S3) associated with the joint phenotype MS-IBD, a subset of which were also significant in cross-trait analyses of MS with UC (N=3; rs2726479, rs116555563, rs67111717) and CD (N=6; rs13428812, rs181826, rs4944014, rs646153, rs10139547, rs11117427). A further 11 and 12 novel SNPs were uniquely associated in joint analyses of MS-UC and MS-CD, respectively, from which only one SNP (e.g. rs1267489 and rs9370774; pairwise *r^2^*=0.92) overlapped. The maxFDR (i.e. the upper bound for the false discovery rate [FDR]) values for MTAG analyses of MS and each of IBD, UC, and CD were roughly 4.55×10^−7^, suggesting our MTAG results were in accordance with the equal variance-covariance assumption.

### Suggestive but inconclusive evidence for causality between MS and UC but not CD

Next, we used bi-directional Mendelian randomization (MR) to explore if genetic overlap between MS and each of IBD, UC and CD was consistent with pleiotropy – as we would intuitively expect – or the presence of causal relationships. We applied multiple (N=6) bi-directional MR methods to each pair of phenotypes (MS-IBD, MS-UC, MS-CD), with the rationale that robust relationships would exhibit consistent and statistically significant results across different methods, including CAUSE (Causal Analysis Using Summary Effect estimates)^26^, which is the only method capable of distinguishing causality from both correlated and uncorrelated pleiotropy. We found consistent evidence for a causal effect of MS on UC and IBD using five of six MR methods (Bonferroni threshold p≤8.33×10^−3^, based on three bi-directional comparisons), but CAUSE could not distinguish a model of causality from correlated pleiotropy for either MS-UC (p=0.16) or MS-IBD (p=0.03; Table S6). In the reverse analyses, there was no or weak and inconsistent evidence for a causal effect of either IBD or UC on MS, and the same was true in bidirectional analyses of MS and CD (Figure 3, Tables S4 & S6). We repeated our analyses with the MHC region excluded, with generally weaker evidence for a causal effect of MS on UC but stronger evidence for a causal effect of MS on CD (Tables S5 & S7, Figure S5).

**Figure 3.**
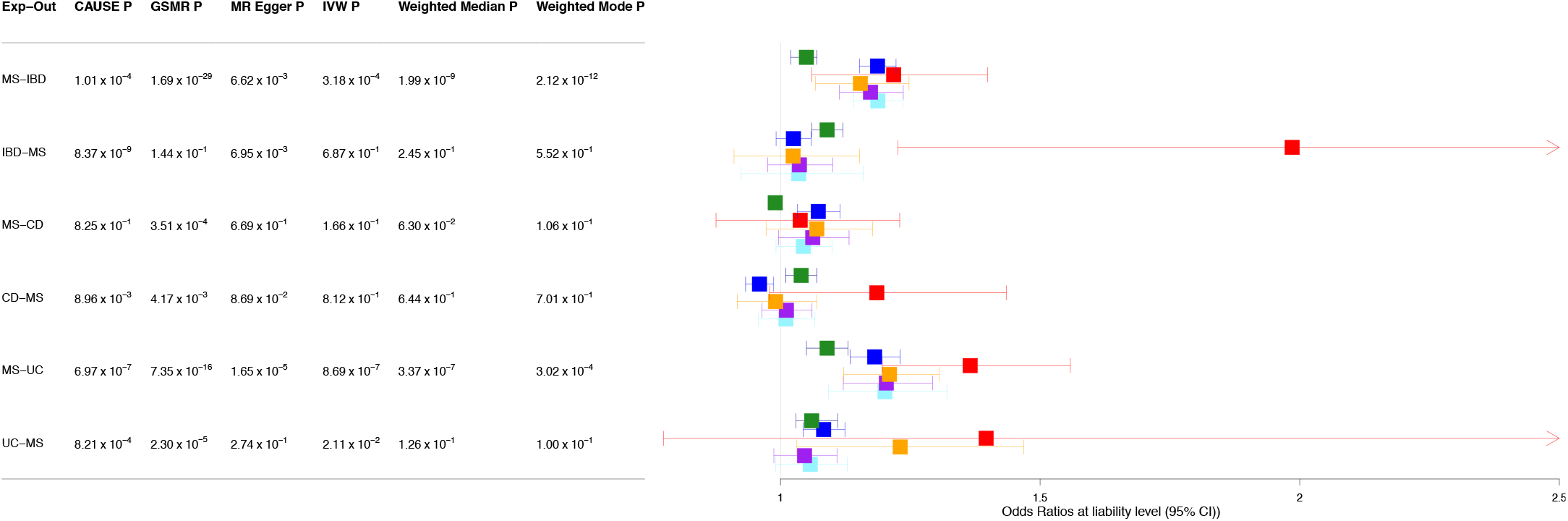
Summary of bi-directional MR analyses between MS and each of IBD, UC and CD. Green: CAUSE; Dark blue: GSMR; red: MR-Egger; orange: IVW; purple: weighted mean; light blue: weighted mode.

### Tissue-level SNP heritability enrichment in MS, IBD, UC and CD

We used stratified LD score regression (S-LDSC)^27^ to evaluate tissue-level enrichment of SNP heritability for MS, IBD, UC and CD. We identified FDR-(p<~5×10^−3^) or Bonferroni-(p<~3×10^−4^) significant SNP heritability enrichment in MS, IBD, UC and CD in lung, spleen, whole blood and small intestine–terminal ileum (with the exception of CD), after adjusting for the baseline model (Figure 4). The magnitude of SNP heritability enrichment in these immune system-related tissues ranged from 2.41 to 3.43 and was largely similar in each disease (Table S9). The enrichment correlations among MS, UC and CD were relatively high and similar for each trait pair, with estimates ranging from 0.80 to 0.85 (see Table S12). Additionally, UC but not MS or CD exhibited Bonferroni-significant enrichment in colon (Figure S8).

**Figure 4.**
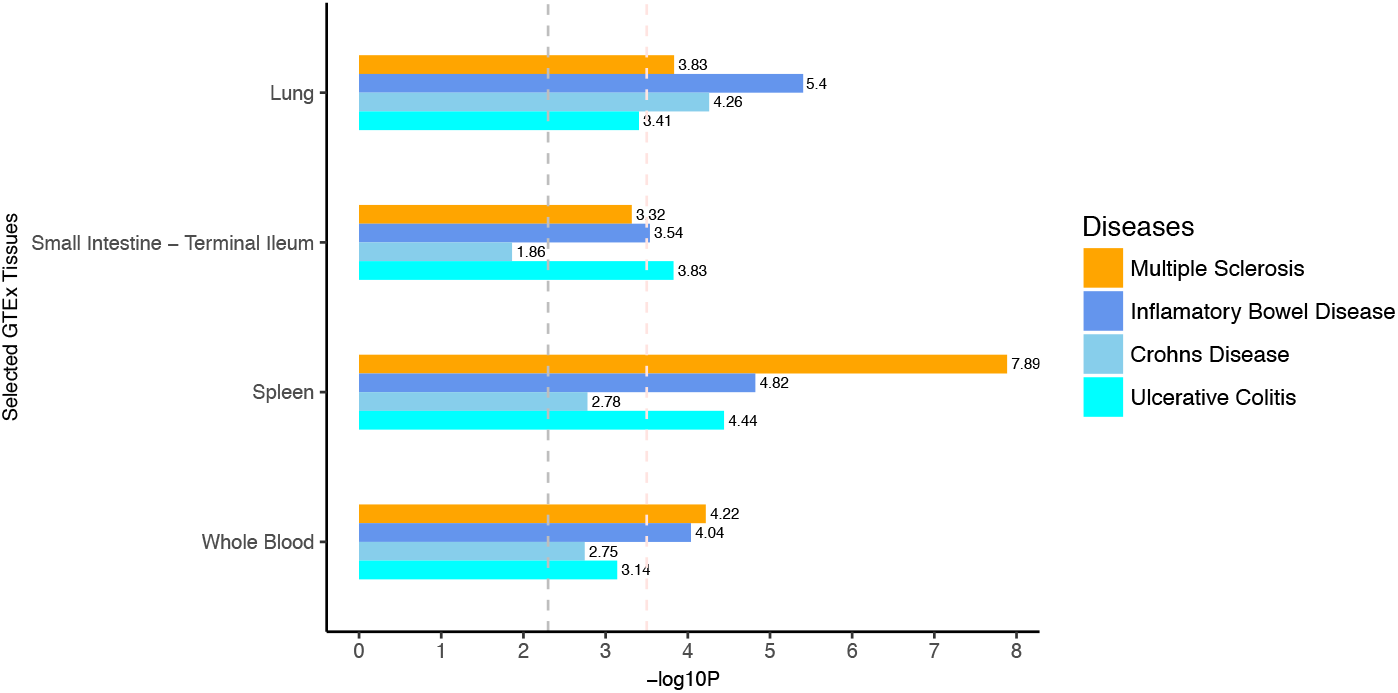
Tissue type-specific enrichment of SNP heritability in MS, IBD, UC and CD in immune tissues. Negative log10 p-values of coefficient Z-scores are displayed on the x axis. The grey and pink dotted lines represent the FDR threshold <5% and Bonferroni corrected threshold, respectively.

### Cell type-level SNP heritability enrichment in MS, IBD, UC and CD

We extended S-LDSC to investigate cell-type specific SNP heritability enrichment for MS, IBD, UC and CD in lung, small intestine–terminal ileum, spleen and peripheral blood. We identified FDR-significant (p<~5×10^−3^) enrichment for all four diseases in CD4^+^ T cells in lung, and enrichment for MS, IBD and CD in CD8^+^ cytotoxic T cells in both lung and spleen, and regulatory T cells in lung (Figure 5, Table S11). SNP heritability enrichment in MS but not other diseases was observed in naïve B cells and dividing T cells in lung, B hypermutation cells in spleen and transitional amplifying cells in small intestine. Conversely, IBD-specific enrichment was identified in CD8^+^ gamma/delta cells in spleen and CD56^+^ natural killer cells in peripheral blood. We also observed enrichment for IBD, UC and/or CD in a number of dendritic cell types in lung, enrichment for CD in dividing natural killer (NK) cells in lung, and enrichment for UC in early enterocytes in small intestine. As summarised in Table S12, the cell-type specific enrichment correlations between MS and each of UC and CD tended to be significantly higher in lung, and were lower and less significant in spleen and small intestine–terminal ileum. The enrichment correlations of MS with all three IBDs in peripheral blood mononuclear cells (PBMC) were similarly estimated at around 0.60, at the marginal significance level. The enrichment correlations between UC and CD persisted to high values at ~0.75 across lung and spleen, which became marginal significant and dropped to ~0.55 in PBMC and small intestine–terminal ileum.

**Figure 5.**
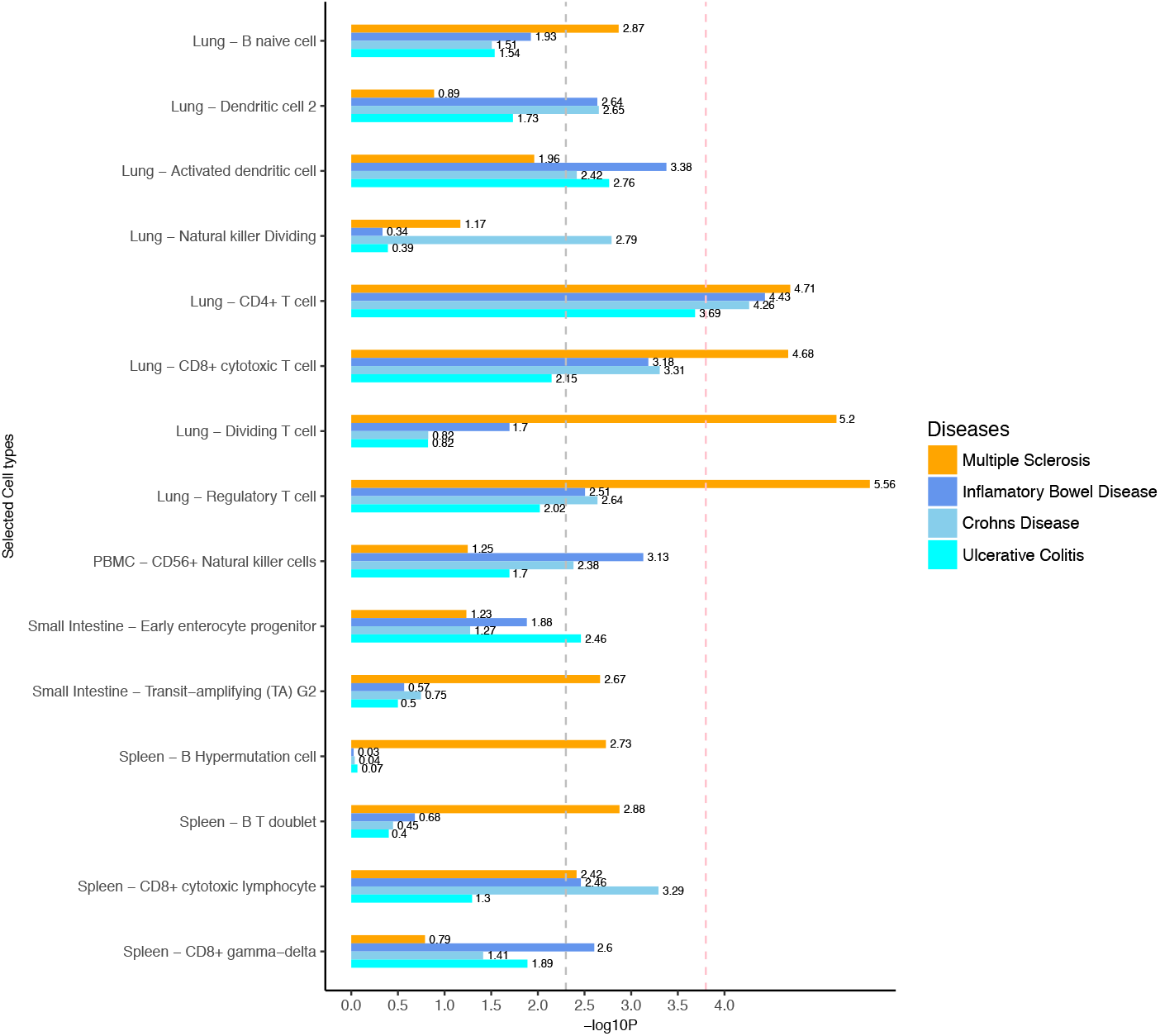
Selected cell type-specific enrichment of SNP heritability in MS, IBD, UC and CD in immune tissues. Cell types are included if they showed FDR significant enrichments in at least one disease. Negative log10 p-values of coefficient Z-scores are displayed on the x axis. The grey and pink dotted line represent the FDR threshold <5% and Bonferroni corrected threshold, respectively.

### Identification of shared functional genes for MS and IBDs

SMR applied to GWAS summary data for MS, IBD, UC and CD and eQTL summary data from eQTLGen (whole blood)^28^ and GTEx (lung, small intestine–terminal ileum, spleen)^29^ identified 210 genome-wide significant associations (p_SMR_<5.36×10^−7^), of which 59 (representing 41 unique genes) survived the HEIDI (HEterogeneity In Dependent Instrument)-outlier test (Table S13, Figure S23-26). Among these 41 genes, the only gene shared by MS and one or more of IBD, UC or CD was *GPR25*, which was significant for MS (p_SMR_=1.18×10^−9^, p_heidi_=0.21), IBD (p_SMR_=2.91×10^−10^, p_HEIDI_=0.12) and UC (p_SMR_=4.98×10^−8^, p_HEIDI_=0.63), but not CD (p_SMR_=4.82×10^−5^, p_HEIDI_=0.63). The remaining genes were associated with either MS (N=25) or one or more of IBD (N=9), UC (N=3) and CD (N=10), with the majority identified in whole blood (N=34) rather than lung (N=9), spleen (N=6) or small intestine–terminal ileum (N=2). Identified genes included three novel genes for MS (antisense gene *LL22NC03-86G7.1*, pseudogene *AC100854.ľ)* and IBD/CD (pseudogene *AL133458.1)*, and numerous previously reported genes for MS (e.g. *CD40*^30^, *MMEL1^31^)* and UC and/or CD (e.g. *CARD9*^32^, *GSDMB*^33^, and *ERAP2^34^).* In addition, we identified 68 significant genes (representing 57 unique genes, Table S14) associated with cross-trait MS-IBD, MS-UC and MS-CD, of which seven passed the HEIDI-outlier test (MS-IBD N=4, MS-UC N=1, MS-CD N=4). Three of these genes were novel *(GDPD3* and two long non-coding RNA [LncRNA] genes *AL031282.2* and *AL109917.1)* and the remaining four (*TNFRSF18*^35^, *DNMT3A^36^,^37^, AHSA2P^38^, NDFIP1*^39,40^) have been previously reported. All seven genes were genome-wide significant using eQTLGen data, but were not identified in the GTEx datasets, probably reflecting lower study power in the latter.

## Discussion

By leveraging large GWAS datasets as well as tissue and cell-type-specific expression data, our study provided novel insights into the shared genetic architecture underlying MS and each of UC and CD.

We identified a stronger genetic correlation between MS and UC than between MS and CD, suggesting that genetic factors make a stronger contribution to comorbidity of MS and UC than MS and CD. Notwithstanding that both genetic and environmental factors may contribute to disease comorbidity, our findings are more consistent with epidemiological reports of stronger comorbidity between MS and UC, as opposed to CD (e.g. Bernstein et al. 2005^7^, Gupta et al. 2015^8^), than with studies reporting no detectable difference in prevalence of MS between patients with UC and CD and vice versa (e.g. Kosmidou et al. 2017^6^). However, we note that Kosmidou et al. 2017^6^ (N=~1 Million) is a much larger study compared to either Bernstein et al. 2005^7^ (N=~8,000) or Gupta et al. 2015^8^(N=~10,000).

Analysis of local genetic correlations between MS and each of IBD, UC and CD were largely compatible with the MR analyses (see below), insomuch as there was no evidence for a causal effect of MS on IBD, UC or CD, or vice versa. The observation that regional *r_g_* estimates were similar to global *r_g_* estimates from LDSC is consistent with the idea that many genetic variants across the genome have pleiotropic effects on these traits. In relation to the MHC region, we observed significant local genetic correlations between MS and UC, but not CD. These results are consistent with prior evidence^18,41,42^ for a stronger shared contribution of the MHC to MS and UC, compared to MS and CD, although the picture is complex because some local genetic correlations for MS-UC were positive whereas others were negative. MR analyses excluding the MHC region were largely consistent with this, with generally weaker evidence for causal effects of MS on UC (and IBD). In addition, we cannot effectively distinguish potentially horizontally pleiotropic SNPs from instrumental (causal) SNPs in the MHC region, because of the complex LD structure in the MHC region.

Cross-trait GWAS meta-analyses identified >40 novel loci shared between MS and IBD, UC and CD. Interestingly, only one of these loci overlapped between MS-UC (rs1267489) and MS-CD (rs9370774; pairwise *r^2^*=0.92) but with different effect directions, consistent with the idea that distinct genetic pathways are shared between MS and UC compared to those between MS and CD. In addition, we found more novel loci associated with MS-CD (N=18) than MS-UC (N=14), and roughly equivalent numbers of novel loci in the MHC region associated with MS-UC (N=3) and MS-CD (N=2). While these results appeared to be opposite to the stronger genetic correlation and local genetic correlations in the MHC regions between MS and UC than between MS and CD, it may be accidentally produced by the complex polygenic architectures shared between MS and UC (and CD).

MR analyses suggested that the genetic correlation between MS and CD is consistent with horizontal pleiotropy, whereas the evidence was inconclusive with respect to the genetic relationship between MS and UC (and IBD). A consistent causal effect of MS on UC (and IBD) was inferred using five of six MR methods, but we could not rule out the possibility of horizontal pleiotropy because CAUSE was unable to distinguish causality from correlated pleiotropy. We note that inference on causality from individual MR methods varied dramatically (e.g. for MS and IBD, GSMR inferred a causal effect of MS on IBD and no effect of IBD on MS, whereas CAUSE inferred a causal effect of IBD on MS), highlighting the importance of considering multiple methods in MR analyses. Larger and more powerful GWAS for MS, and IBDs will be needed to definitively establish (or rule out) the existence of causal relationships between these diseases.

We replicated previous reports of significant SNP heritability enrichment for each of MS^9,19^, IBD^19^, UC^19^, and CD^19^ in multiple immune system-related tissues, including lung, spleen and whole blood. Additionally, we identified heritability enrichment for MS, IBD and UC (but not CD) in small intestine–terminal ileum. We attribute the discovery of this novel tissue-level association to the availability of more powerful GWAS summary statistics. In the case of MS, this is due to larger sample size (i.e. total sample of 41,505 in this study compared to 17,698 for Finucane et al. 2018^19^), whereas for IBD, UC and CD it is due to more sophisticated statistical methods (i.e. earlier GWAS performed using a meta-analysis of 15 cohorts^11^, compared to the individual-level based bivariate linear mixed-effects model with genetic relatedness matrix as random-effects^12^) yielding more genome-wide significant independent SNPs in comparison to earlier GWAS (i.e. 202, 134 and 165 loci for IBD, UC and CD respectively, compared to 110, 23 and 30). Alterations in small intestine physiology have been reported to be responsible for triggering both MS and IBDs. For example, pro-inflammatory T_H_17 (interleukin-17-producing T helper) cells, which are redirected to and regulated by the small intestine^43^, have been implicated in the pathogenesis of both MS^44^ and IBD^45^.

We then extended S-LDSC to the cellular level, identifying a number of novel findings in comparison to Finucane et al. 2018^19^, who performed similar analyses and reported seven cell types (three based on analysis of ImmGen data^46^ - DC.8-4-11b+.MLN [myeloid cells] in mesenteric lymph nodes, T.4.Pa.BDC [T cells] in the pancreas, T.4Mem44h62l.LN [T cells] in subcutaneous lymph nodes - and four cell types – CD4, CD8, B and NK cells of primary blood [including peripheral blood and bone marrow cells] – based on of analysis of haematopoiesis ATAC-seq data^47^) with significant heritability enrichments in MS, IBD and CD, but not UC.

First, we identified SNP heritability enrichments for MS, IBD, UC and CD in CD4^+^ T cells in lung. Several CD4^+^ T cell-related genes have been reported to be involved in risk of MS, UC and CD^48,49^. For instance, *IL23A* (Interleukin-23A), which mediates CD4^+^ T cell function through its receptor IL23R, was reported to be involved in the pathophysiology of MS^50^ and IBD^51^. The *IFNG* gene has also been found to be associated with both MS^52^ and IBD^53^, through regulation of Th1 and Th2 cytokines. Both *IL23A* and *IFNG* were highly expressed in CD4^+^ T cells in lung in our analyses.

Second, we found significant SNP heritability enrichments in CD8^+^ cytotoxic T cells in both lung and spleen as well as regulatory T cells in lung in MS, IBD and CD, but not UC. Interestingly, similar enrichment in these T cells was also observed in UC, but these became non-significant after adjusting for the baseline models, indicating that the enrichment signals from these T cell-specific genes in UC can be explained by pathways associated with the baseline annotations. Several candidate genes were involved in regulation of these T cells and have been implicated in both MS and CD. For example, *PTGER4*, which encodes the prostaglandin receptor, was found to be involved in susceptibility to both MS^54^ and CD^55^, possibly through prostaglandin E2 which is relevant to the immune system via regulation of cytokines^56^. Another gene, *CXCR6*, whose expression is thought to be highly relevant to the immune system via coding of a chemokine receptor protein, has also been reported to be associated with both MS^57^ and CD^58^. Both *PTGER4* and *CXCR6* are highly expressed in these T cells-related genes in our study.

Third, we observed significant heritability enrichment for UC in early enterocytes and for MS in transitional amplifying cells in the small intestine. Epithelial cells in the small intestine are thought to be involved in the pathogenesis of IBD via dysfunction in processing and transmission of antigens to immune cells through the intestinal mucosa ^59^, whereas the role of transitional amplifying cells in MS risks is unknown. We failed to identify any cell types in small intestine showing heritability enrichments in both MS and IBDs, which may be a consequence of insufficient study power and/or reliance on small intestine data from mouse, as opposed to human tissues.

Of note, we did not replicate previously reported SNP heritability enrichments in any PBMC cell type for either MS or IBDs. This observation may be explained by differences in the cell type-specific reference data used in our study (i.e. PBMC data) compared to that in prior papers (e.g. Finucane et al. 2018^19^, bone marrow, haematopoiesis ATAC-seq data^60^). We did replicate significant heritability enrichments in several other cell types, including B cells and NK cells, in either MS or IBDs but not both, indicating the specific pathogenic roles of these cells (compared to CD+ T cells) in triggering MS and IBDs.

We identified several putatively functional genes shared between MS and one or more IBDs. Application of SMR and HEIDI to single-trait GWAS (e.g. MS, IBD) identified three novel genes for MS *(LL22NC03-86G7.1* and *AC100854.1)* or IBD/CD *(AL133458.1)* as well as a single gene *(GPR25)* associated with MS, IBD and UC, which is a G protein-coupled receptor that is highly expressed in T cells and NK cells and has been revealed to be involved in risk of MS and IBD^61^. Moreover, another three novel shared genes *(GDPD3, AL031282.2* and *AL109917.1*) were identified in equivalent analyses of cross-trait GWAS meta-analyses. *GDPD3* has been reported to be implicated in lipid metabolism and adaptive immunity, in particular relating to dendritic cells^62,63^. The other five genes, antisense gene *LL22NC03-86G7.1*, pseudogenes *AC100854.1* and *AL133458.1*,and *AL031282.2* and *AL109917.1* both code for long non-coding RNAs, which as a class of pseudogenes and non-coding RNAs have previously been hypothesized to make crucial contributions to comorbid MS^38,64^ and IBD^65,66^, although the function of these specific genes remains unclear.

Our study had a number of limitations. First, some unmeasured confounding (e.g. history of medication may lead to potential pleiotropic effects that impacts MR effect of exposure on outcome through other pathways modified by medication) may underlie MS and IBD and thus influence the accuracy of MR estimates. However, these effects are likely negligible as we applied multiple MR approaches to minimise the false-positive rate of our results. Secondly, we evaluated the tissue and cell type-specific heritability enrichments on the basis of top 10% most specific genes, which may neglect influences from other genes with less specific effects. Thirdly, we only selected nearby SNPs of the top genes and excluded the SNPs in the MHC region for LD score regression, which may result in underestimation of genetic correlations between MS and IBDs as well as heritability enrichments per tissue and cell for MS and IBDs.

In summary, our study revealed stronger shared genetic variance underlying MS and UC compared to MS and CD, but evidence on whether this represents a causal effect in relation to MS and UC was inconclusive. We identified several novel genetic risk loci and three candidate genes significantly implicated in susceptibility to cross-trait MS and IBD (or UC or CD), none of which was genomewide significant in the single trait (e.g. MS, IBD) GWAS. We revealed evidence for shared SNP heritability enrichment for MS and UC (or IBD) in small intestine–terminal ileum, as well as a group of T cells in lung and/or spleen (i.e. CD4^+^ T cell in lung, CD8^+^ cytotoxic T cell and regulatory T cell in lung and/or spleen), providing further evidence supporting an important contribution of some specific immune-system related tissues and cell types likely enriched in the heritability of MS and IBDs (including the two predominant subtypes UC and CD) and their shared genetic variance. Our findings progress understanding of shared genetic mechanisms underlying MS and IBDs.

## Methods

### Study samples

#### GWAS dataset for MS

GWAS summary results for MS were obtained from the International MS Genetics Consortium (IMSGC) meta-analysis of 15 datasets comprising 14,802 MS cases and 26,703 controls of European ancestry^9^. Each dataset was imputed using the 1000 Genomes European panel. SNPs with minor allele frequency (MAF) >1% were utilised for meta-analysis using a fixed-effects model. As the MAF information was not available in the MS GWAS meta results, we annotated the MAF information based on the European population from the 1000 Genomes panel. Ambiguous SNPs (AT, TA, CG and GC) were excluded and a total of ~6.8 million SNPs were retained for analysis.

#### GWAS datasets for IBD, UC and CD

We obtained publicly available GWAS summary data for UC, CD and IBD, the latter case sample comprising those in both the UC and CD GWAS^12^. We note that UC and CD were the primary focus of our analyses, but we also included IBD, so as to compare the results of our genetic analyses to the epidemiological literature for overlap between MS and IBD, and because the GWAS for IBD has greater power than UC and CD alone. A total of 34,652 participants of European ancestry (12,882 cases and 21,770 controls) were included in the IBD GWAS, from which 27,432 Europeans (6,968 cases and 20,464 controls) and 20,883 Europeans (5,956 cases and 14,927 controls) were included in the UC and CD GWAS, respectively. Nearly 12 million SNPs (~9.5 million with MAF >1%) were included in all three GWAS summary statistics, imputed using the 1000 Genomes Europeans as the reference. Genome-wide association analyses for each disease were conducted using PLINK^67^, adjusted by principal components. More details about the cohorts and quality control (QC) process are explained in Jostins et al. 2012^11^ and Liu et al. 2015^12^.

#### Genotype-Tissue Expression (GTEx) data

GTEx is a public data resource of gene expression in 53 non-diseased human tissues^29^. We used normalised (transcripts per million) GTEx V7 data^68^ to assess tissue type-specific gene expression. After excluding low-quality individuals (N=2, defined as <100 genes with >1 read per million) and genes (N=736, defined as <4 individuals with >1 read per million), we retained data on 53 tissues from a total of 751 individuals, with an average of 220 samples per tissue type. In addition, we also downloaded the GTEx V7 expression quantitative trait locus (eQTL) summary data (see *URLs)* for the downstream analysis.

#### Single-cell RNA sequencing (scRNA-seq) data

On the basis of evidence for tissue-level SNP heritability enrichment in the GTEx analyses, we obtained scRNA-seq unique molecular identifier (UMI) count matrices (see *URLs*) from healthy human lung (N=57,020 cells)^69^, spleen (N=94,257 cells)^69^ and peripheral blood (N=68,579 cells)^70^, and mouse small intestine^71^ (N=7,216 cells). For the latter, we filtered genes with mismatched gene symbols between mouse and human. Procedures for normalisation and quality control of the scRNA-seq data have been described previously^69–71^; we used the cell clustering results reported by the authors. A total of 84 cell types across four tissues were utilised in our study (see Table S8), with an average of 2,703 cells per cell type.

### Statistical analyses

#### LDSC

We used LDSC^23^ to estimate single trait SNP heritabilities for MS, IBD, UC and CD and bivariate LDSC to estimate genetic correlations (*r_g_*) between MS and each of IBD, UC and CD, as well as between UC and CD. We reformatted all GWAS summary statistics to the pre-computed LD scores of the 1000 Genomes Europeans reference. SNPs were excluded if they did not intersect with the reference panel, or if they were located in the MHC region (chromosome 6: 28,477,797-33,448,354), had a MAF <1% or INFO score <0.3. SNP heritability estimates were converted to the liability-scale based on the observed sample prevalence and population prevalence, assuming the latter were 0.3%, 0.4%, 0.29%, and 0.25%^72–74^ for MS, IBD, UC and CD, respectively. Genetic correlation estimates were obtained from the single-trait SNP heritability and cross-trait genetic covariance estimates. We conducted LDSC without constraining the intercept and *r_g_* estimates were considered Bonferroni significant if the p-value was 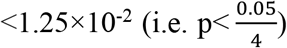. As a sensitivity analysis, we also performed LDSC with the single-trait heritability intercept constrained to evaluate the influence of GWAS statistic inflation.

#### Estimation of local genetic correlations using ρ-HESS

To investigate whether MS shared higher genetic overlap with UC in the local independent genomic region than CD, we applied ρ-HESS^24^ to evaluate the local genetic correlations between MS and each of IBD, UC, and CD. A total of 1,699 default regions that were approximately LD independent with average size of nearly 1.5Mb^75^ were checked by ρ-HESS, including five regions in the MHC (i.e. chromosome 6: 28,017,819–28,917,608, 28,917,608–29,737,971, 30,798,168–31,571,218, 31,571,218–32,682,664, and 32,682,664–33,236,497). We performed ρ-HESS to estimate the local SNP heritability per trait and genetic covariance between traits based on the 1000 Genomes Europeans reference of hg19 genome build. Local genetic correlation estimates were then calculated from the local single-trait SNP heritability and local cross-trait genetic covariance estimates.

#### Multi-Trait Analysis of GWAS

We implemented cross-trait meta-analysis of GWAS summary statistics for MS and each of IBD, UC, and CD, using MTAG^25^. Here, we performed inverse-variance weighted meta-analyses with trait-specific effect sizes that assumes equal SNP heritability for each trait and perfect genetic covariance between traits. We focused on independent genetic variants that were genome-wide significant in the cross-trait meta-analyses (e.g. MS-IBD), but not identified in the original single-trait GWAS (e.g. MS or IBD). These independent genome-wide significant genetic variants were selected using LD clumping *r^2^*<0.05 within 1,000-kb windows through PLINK v1.9^67^ according to the UK Biobank European reference combined imputed by Haplotype Reference Consortium (HRC) and UK10K, a subset of which were excluded if they showed genome-wide significant associations with the original single-trait GWAS of MS or each of IBD, UC, and CD. The upper bound for the false discovery rate (‘maxFDR’) was calculated to examine the assumptions on the equal variance-covariance of shared SNP effect sizes underlying the traits.

#### MR analyses

We used six MR methods to investigate putative causal relationships between MS and each of IBD, UC and CD: Generalised Summary-data-based Mendelian Randomisation (GSMR)^76^, MR-Egger^77^, inverse variance weighting (IVW)^78^, weighted median^79^, weighted mode^80^ and CAUSE^26^. We utilised multiple MR methods with different assumptions on the extent and nature of horizontal pleiotropy, which refers to variants with effects on both outcome and exposure through a pathway other than a causal effect. Horizontal pleiotropy can be correlated, if variants affecting both the outcome and exposure do so via a shared heritable factor, or uncorrelated, if variants affect outcome and exposure traits via separate mechanisms. We considered relationships with consistent evidence for causality using all MR methods to be more reliable and noteworthy.

We used the R packages *GSMR*^76^ and *TwoSampleMR*^7^ to implement five MR methods (GSMR, IVW, MR-Egger, weighted median and weighted mode) with different assumptions about horizontal pleiotropy. Briefly, GSMR assumes no correlated pleiotropy but implements the HEIDI-outlier approach to identify and remove SNPs with evidence for significant uncorrelated pleiotropy. IVW assumes that if uncorrelated pleiotropy is present it has mean zero, so only adding noise to the regression of meta-analysed SNP effects with multiplicative random effects^78^. MR-Egger further allows for the presence of directional (i.e. non-zero mean) uncorrelated pleiotropy and adds an intercept to the IVW regression to exclude confounding from such pleiotropy^77^. Two-sample MR methods capable of accounting for some correlated pleiotropy include the weighted median and the weighted mode. The weighted median measures the weighted median rather than weighted mean of the SNP ratio, which has the ability to identify true causality if ≤50% of the weights are from invalid SNPs^79^. The weighted mode classifies the SNPs into groups according to their estimated causal effects, and assesses evidence for causality using only the largest set of SNPs, which essentially relaxes the assumptions of MR and has the ability to identify the true effect even if a majority of instruments are invalid SNPs^80^. For these five MR methods, independent SNPs (LD clumping *r^2^*<0.05 within 1,000-kb windows using PLINK v1.9^67^, according to the UK Biobank European reference combined imputed by HRC and UK10K) with evidence for genome-wide association (p ≤5×10^−8^) with the ‘exposure’ trait were used as instrumental variables, and merged with the SNPs from the ‘outcome’ trait.

We also used a recently published Bayesian-based MR method called CAUSE that accounts for both correlated and uncorrelated pleiotropy^26^. Compared to the other two-sample MR methods, CAUSE further corrects correlated pleiotropy by evaluating the joint distribution of effect sizes from instrumental SNPs, assuming that the ‘true’ causal effect can influence all instrumental SNPs while the correlated pleiotropy only influences a subset of instrumental SNPs. CAUSE improves the power of MR analysis by including a larger number of LD-pruned SNPs (LD *r^2^*<0.10) with an arbitrary p ≤1×10^−3^ and provides a model comparison approach to distinguish causality from horizontal pleiotropy.

We implemented bi-directional MR analyses using all six methods to investigate the putative causal effect of MS on each of IBD, UC and CD, and vice versa. Due to the complicated LD patterns in the MHC region, here we performed MR analyses with and without SNPs located within the MHC region, to further investigate the effects of MHC region SNPs on putative causal associations between MS and each of IBD, UC and CD. We applied a stricter LD threshold *(r^2^*<0.001) when pruning SNPs in the MHC region.

We declared inferred causal relationships to be significant if they showed Bonferroni-corrected 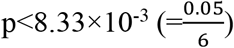 using all MR methods. For all MR methods, we converted our estimated MR effect size from logit-scale to liability-scale using the formula described by Byrne et al. 2019^81^ (i.e. 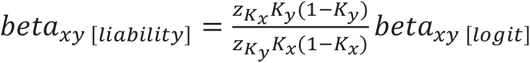, where *K_x_* and *K_y_* are the population prevalence of exposure and outcome trait, respectively; and 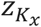 and 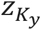 are the height of the Gaussian distribution at the population prevalence threshold for exposure and outcome trait, respectively), assuming the population prevalence for MS, IBD, UC and CD were 0.3%, 0.4%, 0.29%, and 0.25%^72–74^, respectively. We then transformed the liability-scale effect size to an odds ratio.

#### Tissue and cell-type specific enrichment of SNP heritability

##### Selection of tissue type- and cell type-specific expressed genes

We selected genes that were highly expressed in each GTEx tissue and cell type using the method described by Bryois et al. 2020^82^. For GTEx, we followed Bryois et al. in excluding testis and tissues that were non-natural or collected in <100 donors. We then calculated the average gene expression for tissues in the same organ (e.g. colon-sigmoid and colon transverse), with the exception of brain tissues. Subsequently, for each tissue and cell type, we excluded non-protein coding genes, genes with duplicated names, genes located in the MHC region, and genes not expressed in any tissue or cell type. We then scaled gene expression to a total of 1 million UMIs per tissue or cell type, and calculated, for each gene, the proportion (ranging from 0 to 1) of total expression across all tissue/cell types that was specific in each tissue/cell type. The top 10% most highly specific genes for each tissue and cell type were then selected for downstream analyses.

##### Stratified LD score regression

We first used S-LDSC^27^ to investigate whether SNP heritability for MS, IBD, UC and CD was enriched in specific tissues. We then applied S-LDSC to scRNA-seq data to evaluate whether specific cell types in those tissues showed significant heritability enrichment. For each of 37 GTEx tissues and 84 cell types from healthy human lung (N=28), spleen (N=30) and peripheral blood (N=11), and mouse small intestine (N=15; we used mouse small intestine data as a ‘proxy’ because no large human small intestine data is publicly available), we defined a focal functional category by selecting SNPs located within 100Kb (hg19) of the set of 10% most specific genes and added this to the baseline model (comprising 53 genomic annotations). We evaluated the significance of each SNP heritability enrichment estimate using the p-value of the regression coefficient Z-score, after adjusting for the baseline model. Enrichment correlations among MS, UC and CD were calculated by correlating the regression coefficients for GTEx tissues and cell types (by tissues) independently. We adjusted for multiple testing by calculating the Benjamini-Hochberg false discovery rate (FDR), accounting for tissues and cell types separately across the four diseases.

#### Summary-data-based Mendelian Randomisation

We used SMR to identify putative functional genes underlying statistical associations for MS, IBD, UC and CD, as well as novel loci identified in cross-trait meta-analyses of MS-IBD, MS-UC and MS-CD, motivated by the question of whether common risk genes underlie MS and inflammatory bowel diseases. SMR^22^ performs a Mendelian randomisation-equivalent analysis that uses summary statistics from GWAS and eQTL studies to test for an association between gene expression (i.e. exposure) and a target phenotype (i.e. outcome), using genome-wide significant SNPs as instrumental variables. A significant SMR association could be explained by a causal effect (i.e. the causal variant influences disease risk via changes in gene expression), pleiotropy (i.e. the causal variant has pleiotropic effects on gene expression and disease risk) or linkage (i.e. different causal variants exist for gene expression and disease). SMR implements the HEIDI-outlier test to distinguish causality or pleiotropy from linkage, but there is currently no way to distinguish causality from pleiotropy.

We implemented SMR using *cis*-eQTL summary data for whole blood from eQTLgen, a metaanalysis of 14,115 samples^28^, and from GTEx V7^29^ for other significant tissues identified by S-LDSC. We utilised UK Biobank European reference combined imputed by HRC and UK10K to evaluate LD, and only focused on expression probes with eQTL p ≤5×10^−8^. Probes located in the MHC region were ignored because of the complicated LD structure in this region. For MTAG-based cross-trait phenotypes (e.g. MS-IBD), SMR analyses were restricted to novel genetic variants not identified in either of the original single-trait GWAS (e.g. MS or IBD). SMR associations due to causality or pleiotropy were declared significant if they surpassed Bonferroni-correction for the total number of eQTLs analysed (N=93,369, p<5.36×10^−7^) and also passed the HEIDI-outlier test (p>0.05, minimum >10 SNPs.

## Code availability

All code for the analyses are available upon request.

## Acknowledgements

We acknowledge funding support from the Australian National Health and Medical Research Council (JG: 1103418, 1127440) and the Mater Foundation (JG, YY, HM). YZ was supported by an MS Research Australia postdoctoral fellowship as well as NHMRC investigator grant L1 (GNT1173155). We also thank the investigators who have made their datasets freely available to undertake research of this type.

## Author contributions

YY, JG, and YZ designed the study and wrote the manuscript. YY performed the primary analyses, with assistance from BT and YZ (data preparation), HM (S-LDSC), ZZ (MR analyses) and YW (MTAG). JG, BT and YZ supervised the study. All authors contributed to the discussion and revision of the manuscript.

## URLs

CAUSE: https://jean997.github.io/cause/index.html

cis-eQTLGen: https://www.eqtlgen.org/cis-eqtls.html

GTEx eQTL: https://cnsgenomics.com/software/smr/#DataResource

IBDs GWAS: https://www.ebi.ac.uk/gwas/publications/26192919

GSMR: http://cnsgenomics.com/software/gsmr/

GTEx: https://gtexportal.org/home/datasets

GTEx eQTL: https://cnsgenomics.com/software/smr/#DataResource

Human lung and spleen scRNA-seq dataset: https://www.tissuestabilitycellatlas.org/

Human PBMC scRNA-seq dataset: https://support.10xgenomics.com/single-cell-gene-expression/datasets

Mouse small intestine Atlas scRNA-seq dataset: https://singlecell.broadinstitute.org/single_cell/study/SCP44/small-intestinal-epithelium

LD score regression: https://github.com/bulik/ldsc

PLINK: https://www.cog-genomics.org/plink/1.9/

Seurat: https://satijalab.org/seurat/

TwoSampleMR: https://mrcieu.github.io/TwoSampleMR/

## References

1. Filippi, M. et al. Multiple sclerosis. Nat Rev Dis Primers 4, 43 (2018).

2. Baumgart, D.C. & Sandborn, W.J. Inflammatory bowel disease: clinical aspects and established and evolving therapies. Lancet 369, 1641–57 (2007).

3. Alkhawajah, M.M., Caminero, A.B., Freeman, H.J. & Oger, J.J. Multiple sclerosis and inflammatory bowel diseases: what we know and what we would need to know! Mult Scler 19, 259–65 (2013).

4. Minuk, G.Y. & Lewkonia, R.M. Possible familial association of multiple sclerosis and inflammatory bowel disease. N Engl J Med 314, 586 (1986).

5. Kimura, K. et al. Concurrence of inflammatory bowel disease and multiple sclerosis. Mayo Clin Proc 75, 802–6 (2000).

6. Kosmidou, M. et al. Multiple sclerosis and inflammatory bowel diseases: a systematic review and meta-analysis. J Neurol 264, 254–259 (2017).

7. Bernstein, C.N., Wajda, A. & Blanchard, J.F. The clustering of other chronic inflammatory diseases in inflammatory bowel disease: a population-based study. Gastroenterology 129, 827–36 (2005).

8. Gupta, G., Gelfand, J.M. & Lewis, J.D. Increased risk for demyelinating diseases in patients with inflammatory bowel disease. Gastroenterology 129, 819–26 (2005).

9. International Multiple Sclerosis Genetics Consortium. Multiple sclerosis genomic map implicates peripheral immune cells and microglia in susceptibility. Science 365 (2019).

10. Chen, G.B. et al. Estimation and partitioning of (co)heritability of inflammatory bowel disease from GWAS and immunochip data. Hum Mol Genet 23, 4710–20 (2014).

11. Jostins, L. et al. Host-microbe interactions have shaped the genetic architecture of inflammatory bowel disease. Nature 491, 119–24 (2012).

12. Liu, J.Z. et al. Association analyses identify 38 susceptibility loci for inflammatory bowel disease and highlight shared genetic risk across populations. Nat Genet 47, 979–986 (2015).

13. International Multiple Sclerosis Genetics, C. et al. Risk alleles for multiple sclerosis identified by a genomewide study. N Engl J Med 357, 851–62 (2007).

14. Belarif, L. et al. IL-7 receptor influences anti-TNF responsiveness and T cell gut homing in inflammatory bowel disease. J Clin Invest 129, 1910–1925 (2019).

15. de Lange, K.M. & Barrett, J.C. Understanding inflammatory bowel disease via immunogenetics. J Autoimmun 64, 91–100 (2015).

16. Restrepo, N.A., Butkiewicz, M., McGrath, J.A. & Crawford, D.C. Shared Genetic Etiology of Autoimmune Diseases in Patients from a Biorepository Linked to De-identified Electronic Health Records. Front Genet 7, 185 (2016).

17. Richard-Miceli, C. & Criswell, L.A. Emerging patterns of genetic overlap across autoimmune disorders. Genome Med 4, 6 (2012).

18. Fernando, M.M. et al. Defining the role of the MHC in autoimmunity: a review and pooled analysis. PLoS Genet 4, e1000024 (2008).

19. Finucane, H.K. et al. Heritability enrichment of specifically expressed genes identifies disease-relevant tissues and cell types. Nat Genet 50, 621–629 (2018).

20. Rodrigues, S. et al. Case series: ulcerative colitis, multiple sclerosis, and interferon-beta 1a. Inflamm Bowel Dis 16, 2001–3 (2010).

21. Kaltsonoudis, E., Voulgari, P.V., Konitsiotis, S. & Drosos, A.A. Demyelination and other neurological adverse events after anti-TNF therapy. Autoimmun Rev 13, 54–8 (2014).

22. Zhu, Z. et al. Integration of summary data from GWAS and eQTL studies predicts complex trait gene targets. Nat Genet 48, 481–7 (2016).

23. Bulik-Sullivan, B.K. et al. LD Score regression distinguishes confounding from polygenicity in genome-wide association studies. Nat Genet 47, 291–5 (2015).

24. Shi, H., Mancuso, N., Spendlove, S. & Pasaniuc, B. Local Genetic Correlation Gives Insights into the Shared Genetic Architecture of Complex Traits. Am J Hum Genet 101, 737–751 (2017).

25. Turley, P. et al. Multi-trait analysis of genome-wide association summary statistics using MTAG. Nat Genet 50, 229–237 (2018).

26. Morrison, J., Knoblauch, N., Marcus, J.H., Stephens, M. & He, X. Mendelian randomization accounting for correlated and uncorrelated pleiotropic effects using genome-wide summary statistics. Nat Genet 52, 740–747 (2020).

27. Finucane, H.K. et al. Partitioning heritability by functional annotation using genome-wide association summary statistics. Nat Genet 47, 1228–35 (2015).

28. Võsa, U. et al. Unraveling the polygenic architecture of complex traits using blood eQTL metaanalysis. bioRxiv, 447367 (2018).

29. GTEx Consortium et al. Genetic effects on gene expression across human tissues. Nature 550, 204–213 (2017).

30. Aarts, S. et al. The CD40-CD40L Dyad in Experimental Autoimmune Encephalomyelitis and Multiple Sclerosis. Front Immunol 8, 1791 (2017).

31. Ban, M. et al. A non-synonymous SNP within membrane metalloendopeptidase-like 1 (MMEL1) is associated with multiple sclerosis. Genes Immun 11, 660–4 (2010).

32. Lamas, B. et al. CARD9 impacts colitis by altering gut microbiota metabolism of tryptophan into aryl hydrocarbon receptor ligands. Nat Med 22, 598–605 (2016).

33. Soderman, J., Berglind, L. & Almer, S. Gene Expression-Genotype Analysis Implicates GSDMA, GSDMB, and LRRC3C as Contributors to Inflammatory Bowel Disease Susceptibility. Biomed Res Int 2015, 834805 (2015).

34. Zerenturk, E.J., Sharpe, L.J. & Brown, A.J. DHCR24 associates strongly with the endoplasmic reticulum beyond predicted membrane domains: implications for the activities of this multi-functional enzyme. Biosci Rep 34 (2014).

35. Croft, M. et al. TNF superfamily in inflammatory disease: translating basic insights. Trends Immunol 33, 144–52 (2012).

36. Low, D., Mizoguchi, A. & Mizoguchi, E. DNA methylation in inflammatory bowel disease and beyond. World J Gastroenterol 19, 5238–49 (2013).

37. Celarain, N. & Tomas-Roig, J. Aberrant DNA methylation profile exacerbates inflammation and neurodegeneration in multiple sclerosis patients. J Neuroinflammation 17, 21 (2020).

38. James, T. et al. Impact of genetic risk loci for multiple sclerosis on expression of proximal genes in patients. Hum Mol Genet 27, 912–928 (2018).

39. Ramon, H.E. et al. The ubiquitin ligase adaptor Ndfip1 regulates T cell-mediated gastrointestinal inflammation and inflammatory bowel disease susceptibility. Mucosal Immunol 4, 314–24 (2011).

40. Altin, J.A. et al. Ndfip1 mediates peripheral tolerance to self and exogenous antigen by inducing cell cycle exit in responding CD4+ T cells. Proc Natl Acad Sci U S A 111, 2067–74 (2014).

41. Goyette, P. et al. High-density mapping of the MHC identifies a shared role for HLA-DRB1*01:03 in inflammatory bowel diseases and heterozygous advantage in ulcerative colitis. Nat Genet 47, 172–9 (2015).

42. International Multiple Sclerosis Genetics Consortium et al. Genetic risk and a primary role for cell-mediated immune mechanisms in multiple sclerosis. Nature 476, 214–9 (2011).

43. Esplugues, E. et al. Control of TH17 cells occurs in the small intestine. Nature 475, 514–8 (2011).

44. Camara-Lemarroy, C.R., Metz, L., Meddings, J.B., Sharkey, K.A. & Wee Yong, V. The intestinal barrier in multiple sclerosis: implications for pathophysiology and therapeutics. Brain 141, 1900–1916 (2018).

45. Monteleone, I., Pallone, F. & Monteleone, G. Th17-related cytokines: new players in the control of chronic intestinal inflammation. BMC Med 9, 122 (2011).

46. Heng, T.S., Painter, M.W. & Immunological Genome Project, C. The Immunological Genome Project: networks of gene expression in immune cells. Nat Immunol 9, 1091–4 (2008).

47. Corces, M.R. et al. Lineage-specific and single-cell chromatin accessibility charts human hematopoiesis and leukemia evolution. Nat Genet 48, 1193–203 (2016).

48. Kamikozuru, K. et al. The expression profile of functional regulatory T cells, CD4+CD25high+/forkhead box protein P3+, in patients with ulcerative colitis during active and quiescent disease. Clin Exp Immunol 156, 320–7 (2009).

49. Peeters, L.M. et al. Cytotoxic CD4+ T Cells Drive Multiple Sclerosis Progression. Front Immunol 8, 1160 (2017).

50. Li, F.F. et al. Characterization of variations in IL23A and IL23R genes: possible roles in multiple sclerosis and other neuroinflammatory demyelinating diseases. Aging (Albany NY) 8, 2734–2746 (2016).

51. McGovern, D. & Powrie, F. The IL23 axis plays a key role in the pathogenesis of IBD. Gut 56, 1333–6 (2007).

52. Kantarci, O.H. et al. IFNG polymorphisms are associated with gender differences in susceptibility to multiple sclerosis. Genes Immun 6, 153–61 (2005).

53. Gonsky, R. et al. IFNG rs1861494 polymorphism is associated with IBD disease severity and functional changes in both IFNG methylation and protein secretion. Inflamm Bowel Dis 20, 1794–801 (2014).

54. De Jager, P.L. et al. Meta-analysis of genome scans and replication identify CD6, IRF8 and TNFRSF1A as new multiple sclerosis susceptibility loci. Nat Genet 41, 776–82 (2009).

55. Glas, J. et al. PTGER4 expression-modulating polymorphisms in the 5p13.1 region predispose to Crohn’s disease and affect NF-kappaB and XBP1 binding sites. PLoS One 7, e52873 (2012).

56. Sander, W.J., O’Neill, H.G. & Pohl, C.H. Prostaglandin E2 As a Modulator of Viral Infections. Front Physiol 8, 89 (2017).

57. Hoglund, R.A. & Maghazachi, A.A. Multiple sclerosis and the role of immune cells. World J Exp Med 4, 27–37 (2014).

58. Diegelmann, J. et al. Expression and regulation of the chemokine CXCL16 in Crohn’s disease and models of intestinal inflammation. Inflamm Bowel Dis 16, 1871–81 (2010).

59. Roda, G. et al. Intestinal epithelial cells in inflammatory bowel diseases. World J Gastroenterol 16, 4264–71 (2010).

60. Buenrostro, J.D. et al. Single-cell chromatin accessibility reveals principles of regulatory variation. Nature 523, 486–90 (2015).

61. Ricano-Ponce, I. et al. Refined mapping of autoimmune disease associated genetic variants with gene expression suggests an important role for non-coding RNAs. J Autoimmun 68, 6274 (2016).

62. Eisenbarth, S.C. et al. NLRP10 is a NOD-like receptor essential to initiate adaptive immunity by dendritic cells. Nature 484, 510–3 (2012).

63. Kaji, T. et al. CD4 memory T cells develop and acquire functional competence by sequential cognate interactions and stepwise gene regulation. Int Immunol 28, 267–82 (2016).

64. Yang, X., Wu, Y., Zhang, B. & Ni, B. Noncoding RNAs in multiple sclerosis. Clin Epigenetics 10, 149 (2018).

65. Lin, L. et al. Which long noncoding RNAs and circular RNAs contribute to inflammatory bowel disease? Cell Death Dis 11, 456 (2020).

66. Harbord, M., Hankin, A., Bloom, S. & Mitchison, H. Association between p47phox pseudogenes and inflammatory bowel disease. Blood 101, 3337 (2003).

67. Purcell, S. et al. PLINK: a tool set for whole-genome association and population-based linkage analyses. Am J Hum Genet 81, 559–75 (2007).

68. GTEx Consortium. The Genotype-Tissue Expression (GTEx) project. Nat Genet 45, 580–5 (2013).

69. Madissoon, E. et al. scRNA-seq assessment of the human lung, spleen, and esophagus tissue stability after cold preservation. Genome Biol 21, 1 (2019).

70. Zheng, G.X. et al. Massively parallel digital transcriptional profiling of single cells. Nat Commun 8, 14049 (2017).

71. Haber, A.L. et al. A single-cell survey of the small intestinal epithelium. Nature 551, 333–339 (2017).

72. Shivashankar, R., Tremaine, W.J., Harmsen, W.S. & Loftus, E.V., Jr. Incidence and Prevalence of Crohn’s Disease and Ulcerative Colitis in Olmsted County, Minnesota From 1970 Through 2010. Clin Gastroenterol Hepatol 15, 857–863 (2017).

73. Hemani, G. et al. The MR-Base platform supports systematic causal inference across the human phenome. Elife 7 (2018).

74. Wallin, M.T. et al. The prevalence of MS in the United States: A population-based estimate using health claims data. Neurology 92, e1029–1040 (2019).

75. Berisa, T. & Pickrell, J.K. Approximately independent linkage disequilibrium blocks in human populations. Bioinformatics 32, 283–5 (2016).

76. Zhu, Z. et al. Causal associations between risk factors and common diseases inferred from GWAS summary data. Nat Commun 9, 224 (2018).

77. Burgess, S. & Thompson, S.G. Interpreting findings from Mendelian randomization using the MR-Egger method. Eur J Epidemiol 32, 377–389 (2017).

78. Burgess, S., Butterworth, A. & Thompson, S.G. Mendelian randomization analysis with multiple genetic variants using summarized data. Genet Epidemiol 37, 658–65 (2013).

79. Bowden, J., Davey Smith, G., Haycock, P.C. & Burgess, S. Consistent Estimation in Mendelian Randomization with Some Invalid Instruments Using a Weighted Median Estimator. Genet Epidemiol 40, 304–14 (2016).

80. Hartwig, F.P., Davey Smith, G. & Bowden, J. Robust inference in summary data Mendelian randomization via the zero modal pleiotropy assumption. Int J Epidemiol 46, 1985–1998 (2017).

81. Byrne, E.M. et al. Conditional GWAS analysis to identify disorder-specific SNPs for psychiatric disorders. Mol Psychiatry (2020).

82. Bryois, J. et al. Genetic identification of cell types underlying brain complex traits yields insights into the etiology of Parkinson’s disease. Nat Genet 52, 482–493 (2020).

