## Supplemental Figures for "Investigating the shared genetic architecture between multiple sclerosis and inflammatory bowel diseases"

### Supplementary Note

#### Heritability enrichments of GTEx tissues using Finucane et al (2018) method

For sensitivity analyses, we applied Finucane et al. 2018[1] method to select the top expressed genes per tissue and performed stratified LD score regression using such gene list per tissue. Again, we excluded tissues that were non-natural tissues, testis tissue, and collected with <100 samples. Here we did not combine the tissues from the same organ and a total of 45 GTEx tissues remained. We then calculated T-statistics per gene for a focal tissue as a proxy, to measure the expression levels in this focal tissue compared to the expressions in all other GTEx tissues ('control' tissues), excepting the tissues from the same tissue region type (e.g., brain-related tissues were excluded from 'control' tissues if the focal tissue was another brain tissue). T-statistics were calculated using linear models via ordinary least-squares. Tissue region type were re-coded to '1' for the focal tissue and '-1' for all the 'control' tissues, and used as a covariate in the linear model. Both age and sex were also included as covariates. Only protein-coding genes were analysed in our study. The top 10% highly expressed genes for each GTEx tissue were then selected for downstream analyses by ranking the T-statistics. In results (Figure S10, Table S10), we observed FDR significant heritability enrichments in both MS and at least one IBDs from lung, small intestine–terminal ileum, spleen, and whole blood, which are close to what we found using Bryois et al. 2020[2] method (Figure S8, Table S9).

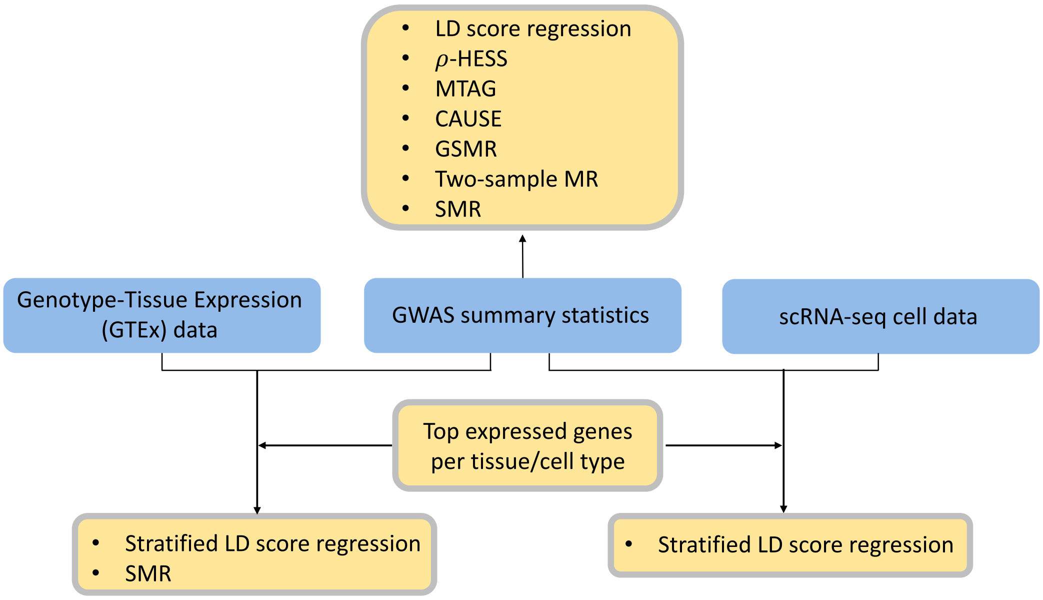

**Figure S1.** Outline of statistical analyses performed in the study.

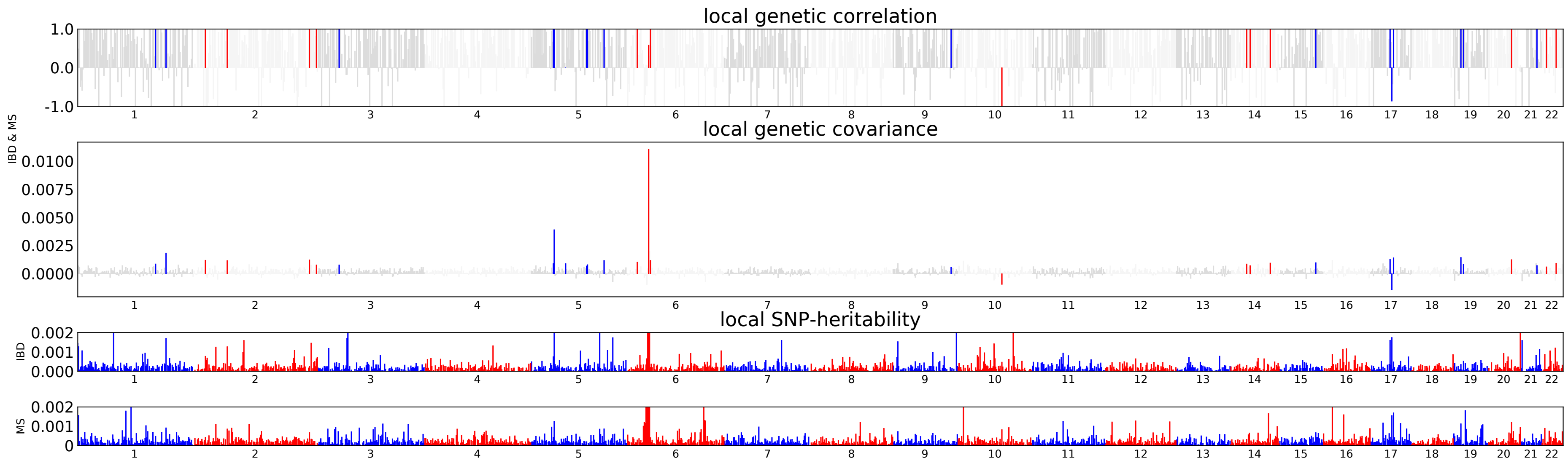

**Figure S2.** Local genetic correlations between MS and IBD revealed by ρ-HESS. Significant local $r_g$ is highlighted in colour red or blue for even or odd chromosomes.

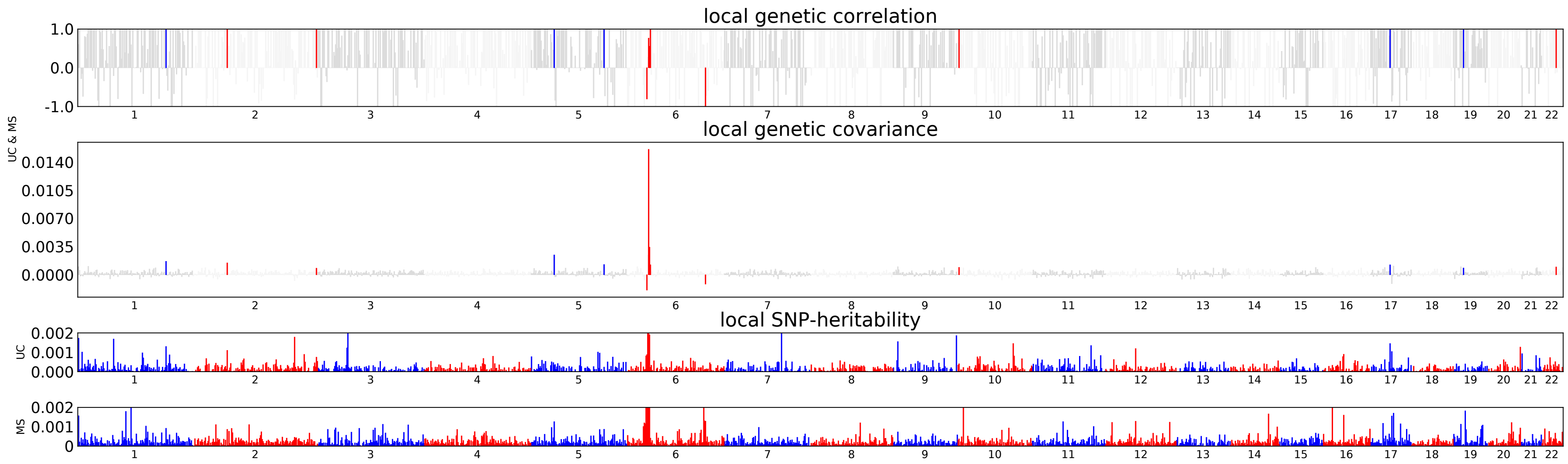

**Figure S3.** Local genetic correlations between MS and UC revealed by ρ-HESS. Significant local $r_g$ is highlighted in colour red or blue for even or odd chromosomes.

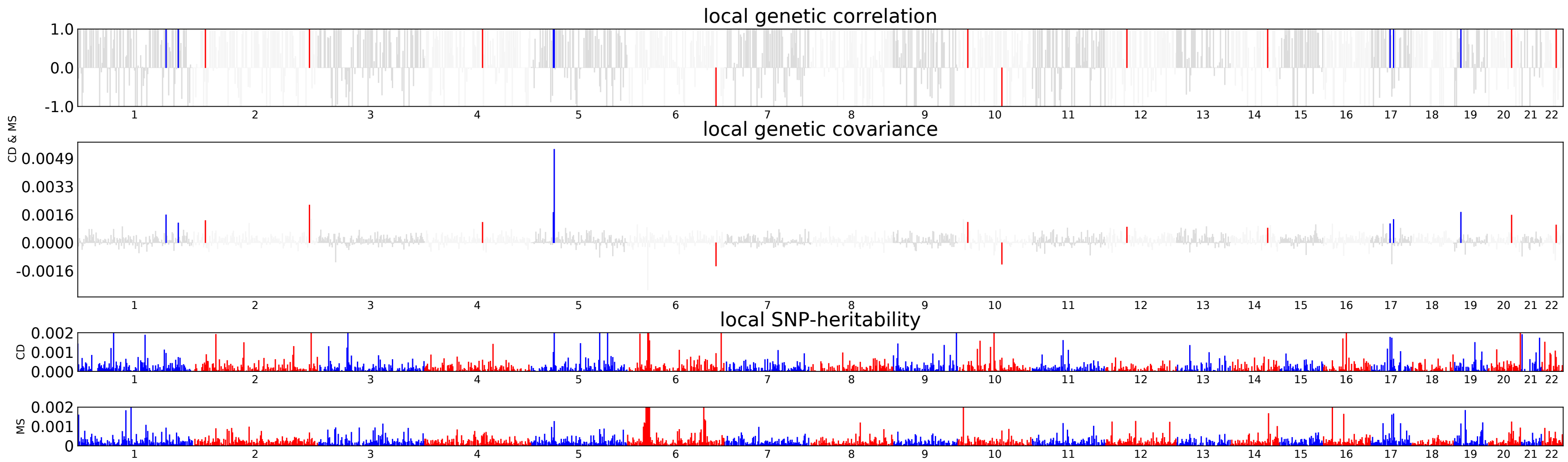

**Figure S4.** Local genetic correlations between MS and CD revealed by ρ-HESS. Significant local $r_g$ is highlighted in colour red or blue for even or odd chromosomes.

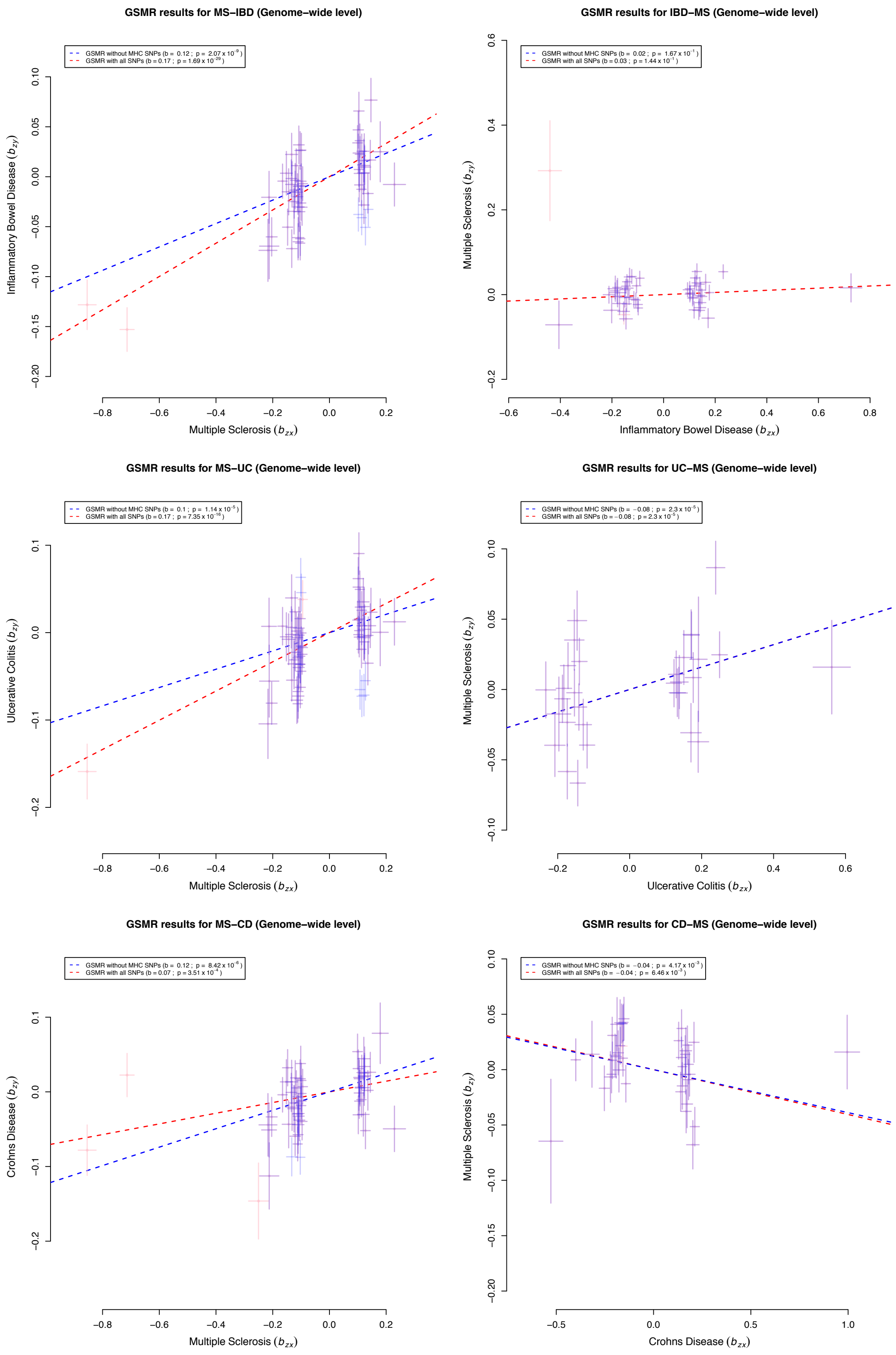

**Figure S5.** Forward and reverse GSMR results of effect sizes (with and without MHC SNPs) for the associations between MS and each of IBD, UC and CD at genome-wide level. For each plot, dots in pink represents the MHC SNPs; dots in purple represents non-MHC SNPs used in either GSMR with or without excluding MHC SNPs; and dots in blue represents the SNPs included by HEIDI for the GSMR without MHC SNPs but removed by HEIDI for the GSMR with all SNPs.

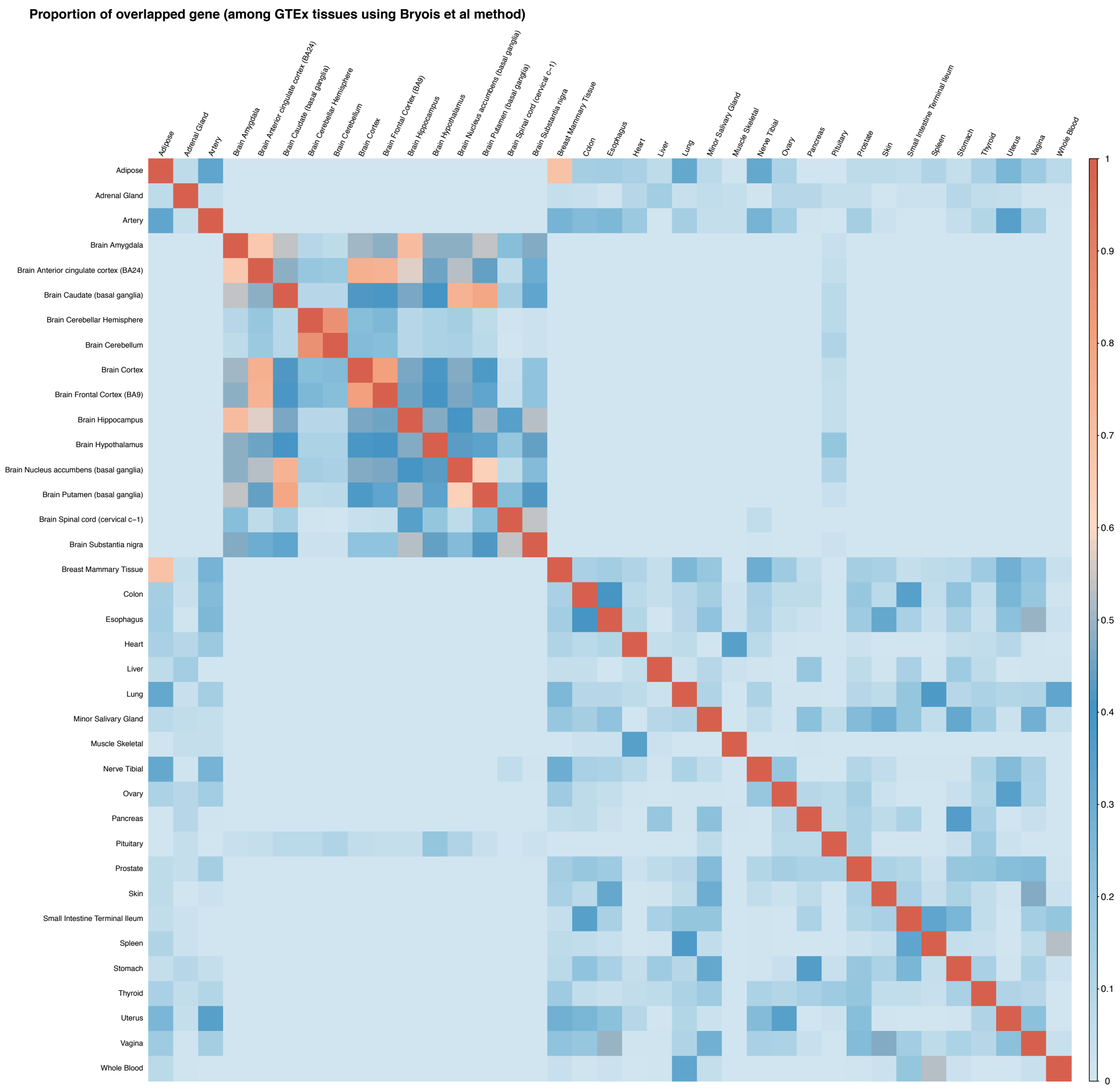

**Figure S6.** Proportion of overlapped top 10% highly expressed genes among 37 GTEx tissues using Bryois et al. 2020 method.

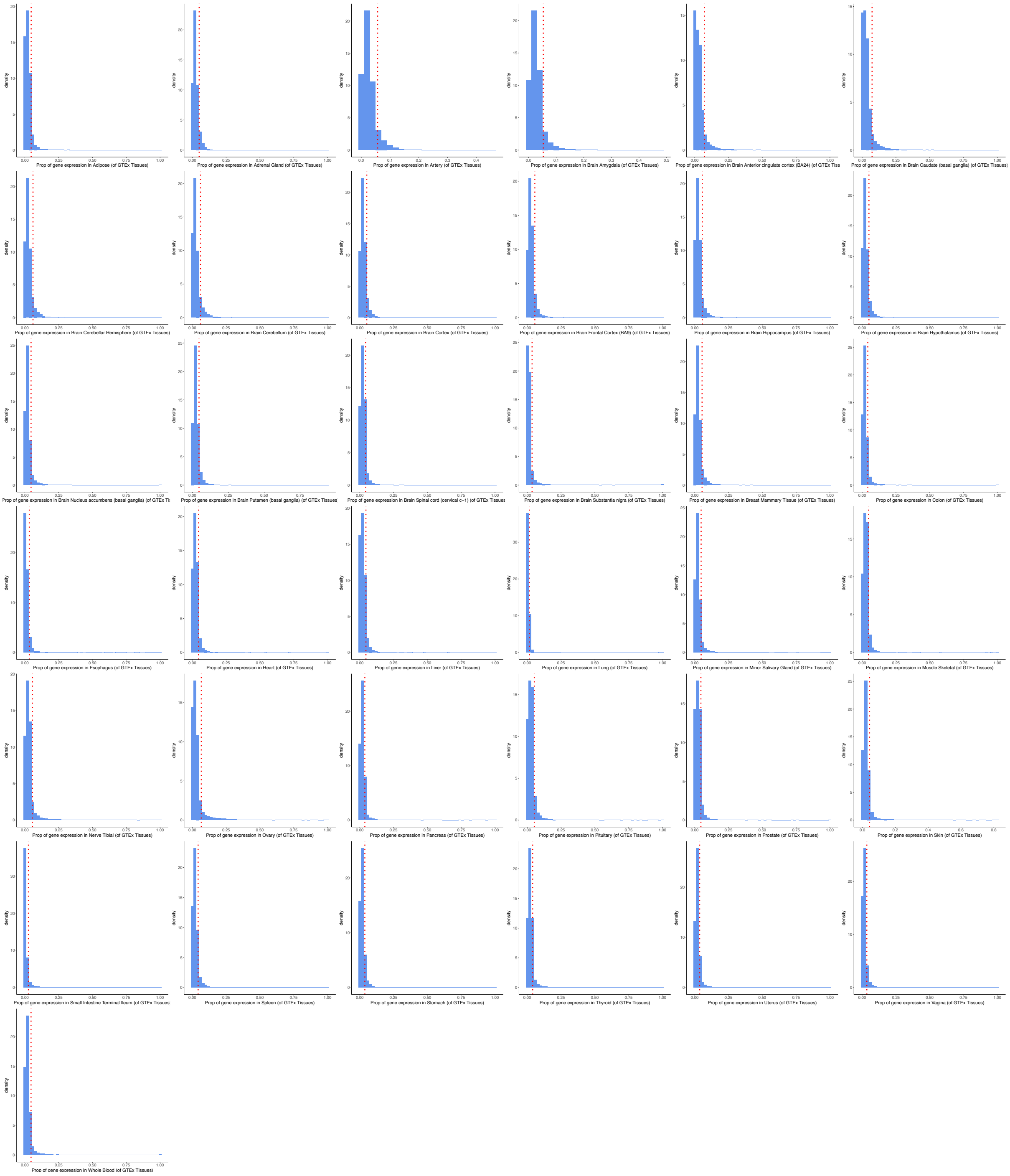

**Figure S7.** Distribution of the proportion of gene expression per GTEx tissue in the total gene expressions of all 37 GTEx tissues applied using Bryois et al. 2020 method. The top 10% expressed genes are distributed in the right parts of the red dotted vertical line.

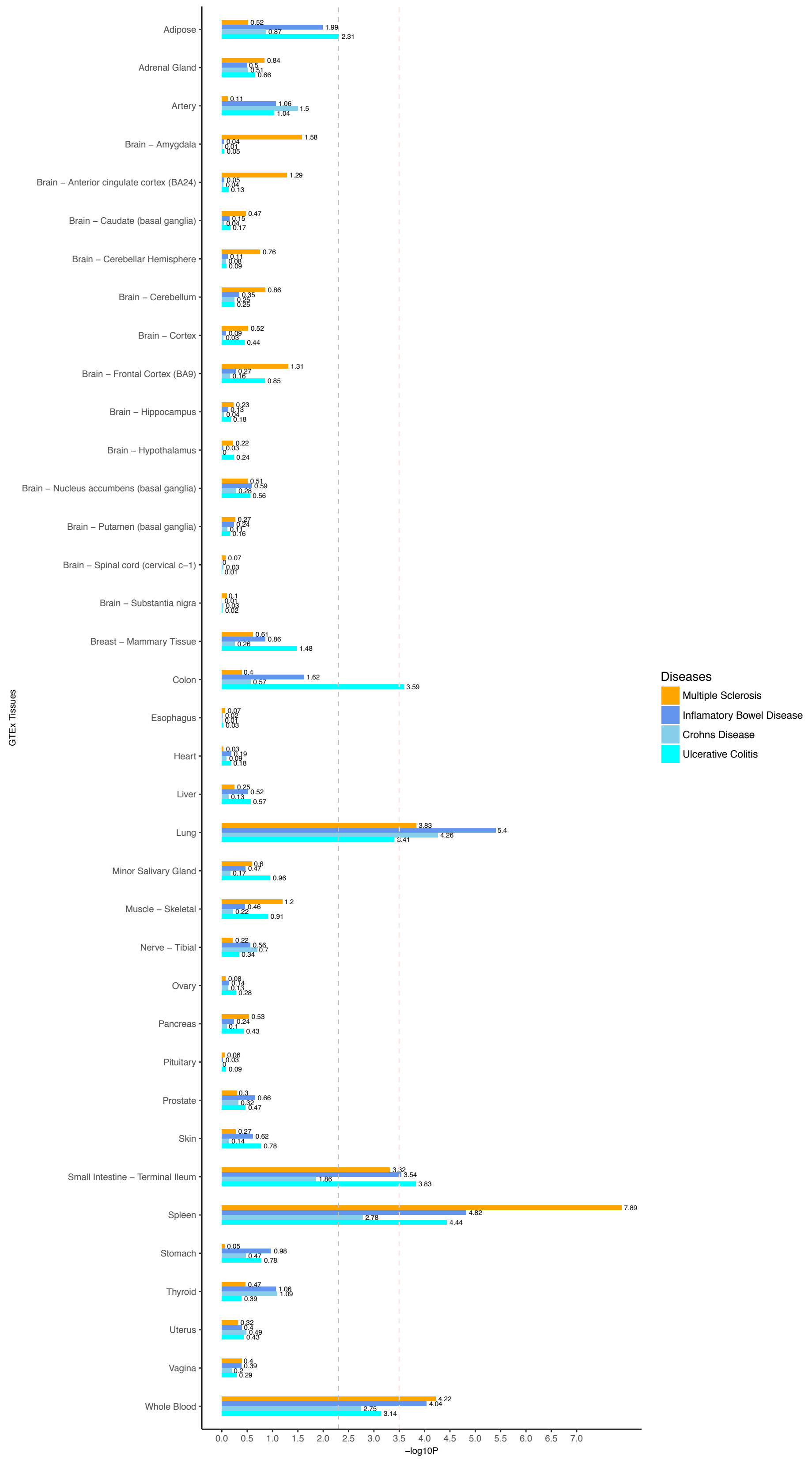

**Figure S8.** Heritability enrichment of 37 GTEx tissue-specific expressed genes in MS and each of IBD, UC, and CD, using Bryois et al. 2020 method. Negative log10 p-value of coefficient Z-score are displayed in x axis. The grey and pink dotted line represent the FDR threshold <5% and Bonferroni corrected threshold, respectively.

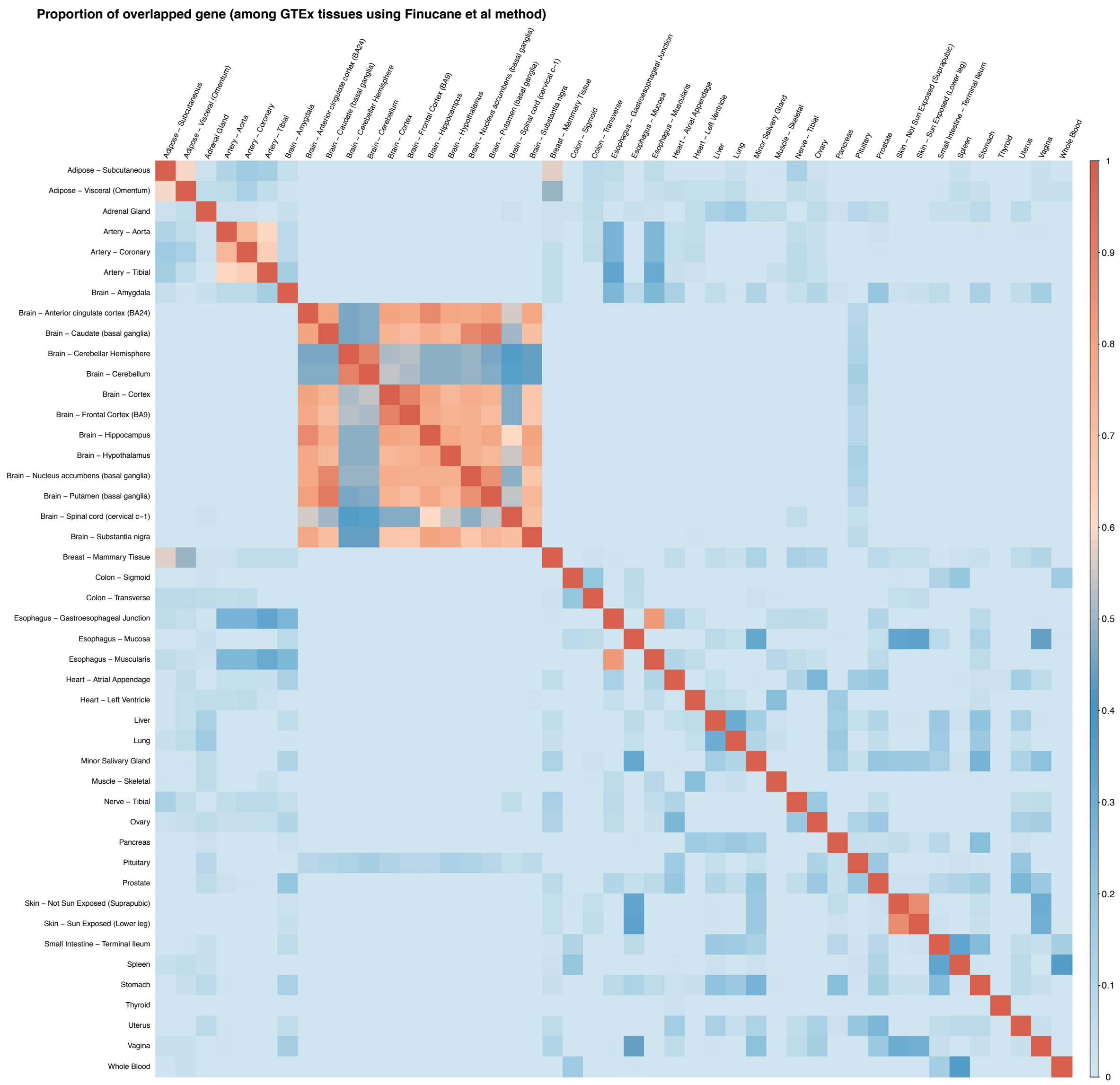

**Figure S9.** Proportion of overlapped top 10% highly expressed genes among 45 GTEx tissues using Finucane et al. 2018 method.

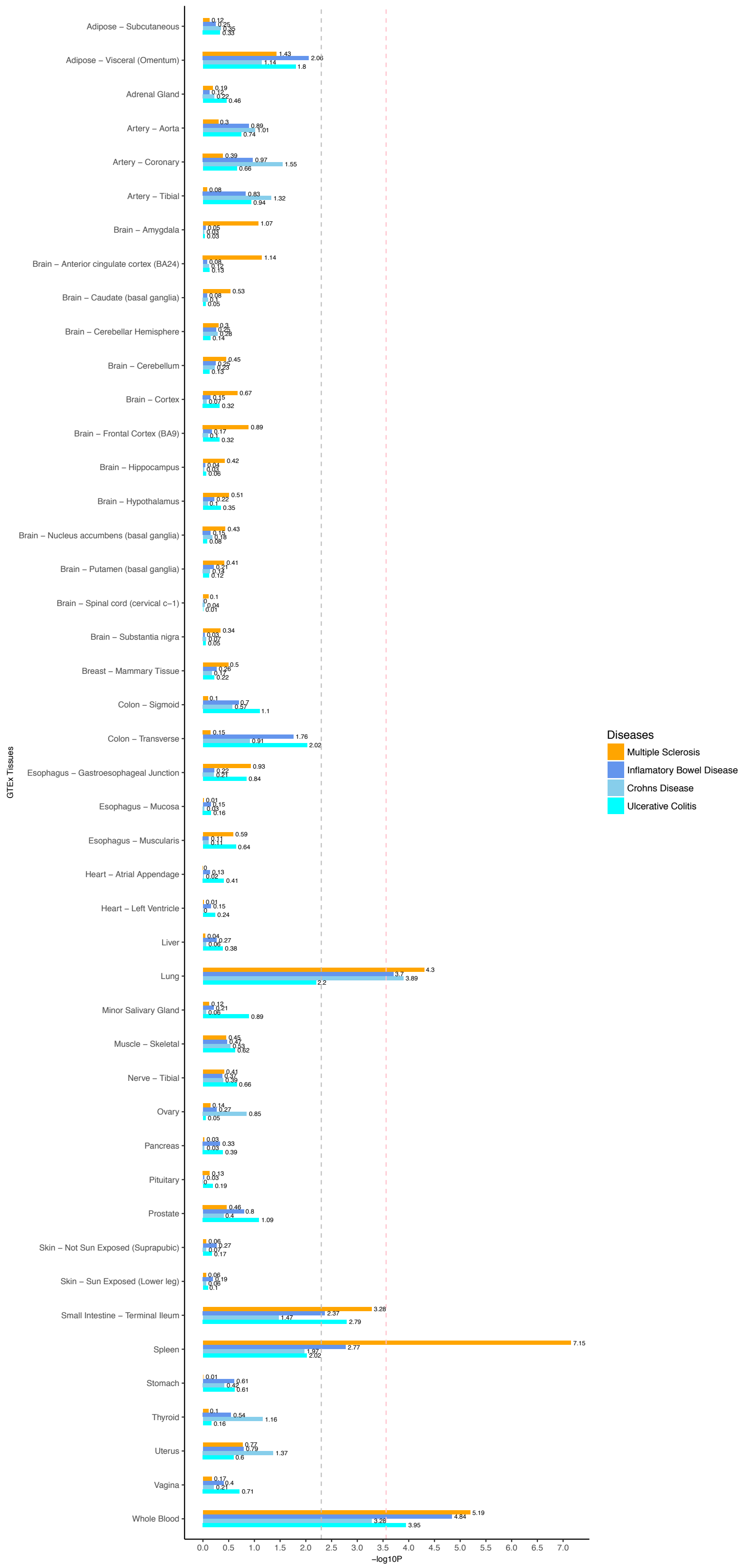

**Figure S10.** Heritability enrichment of 45 GTEx tissue-specific expressed genes in MS and each of IBD, UC, and CD, using Finucane et al. 2018 method. Negative log10 p-value of coefficient Z-score are displayed in x axis. The grey and pink dotted line represent the FDR threshold <5% and Bonferroni corrected threshold, respectively.

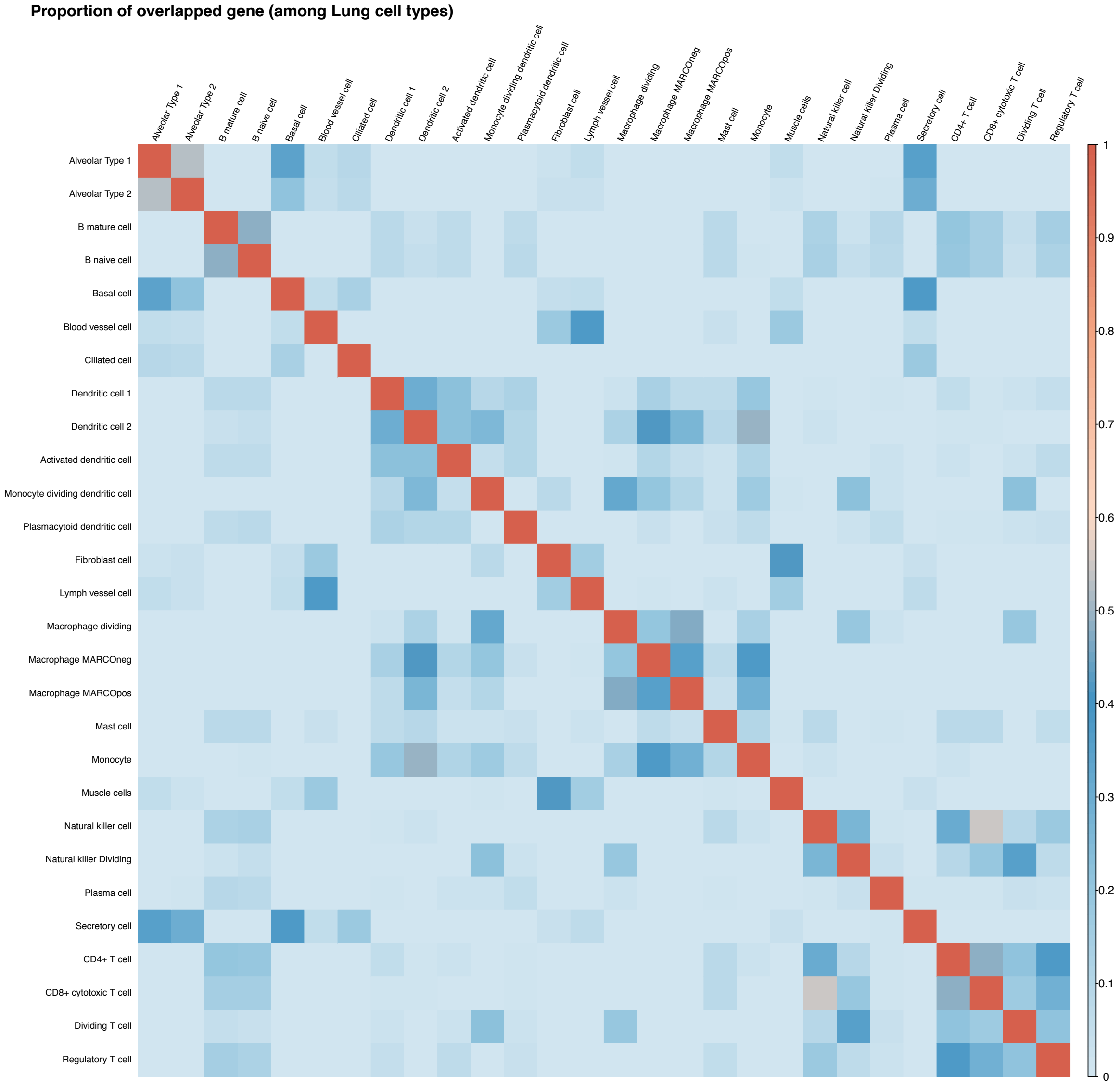

**Figure S11.** Proportion of overlapped top 10% highly expressed genes among 28 lung cell types.

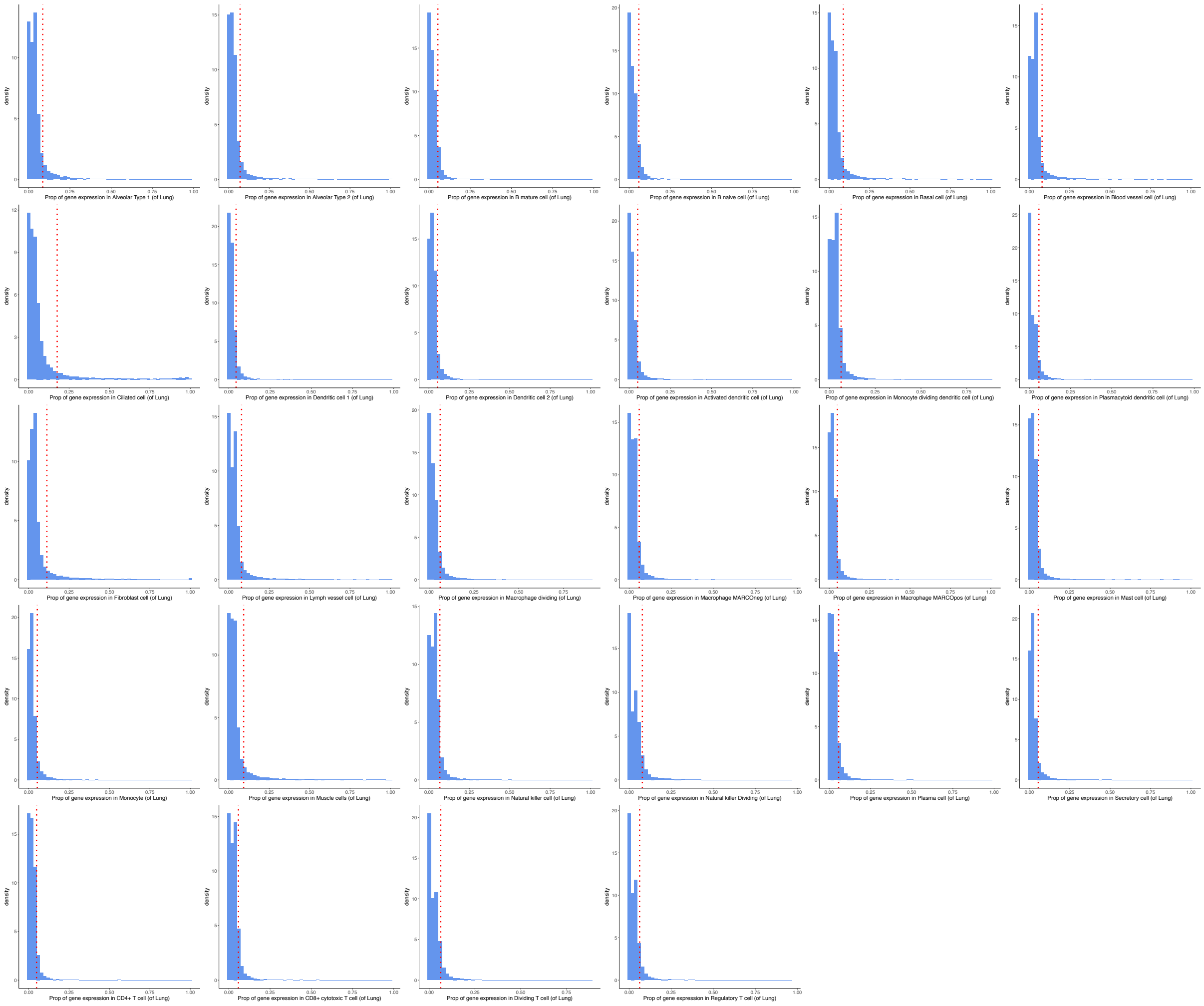

**Figure S12.** Distribution of the proportion of gene expression per cell type in the total gene expressions of all 28 lung cell types. The top 10% expressed genes are distributed in the right parts of the red dotted vertical line.

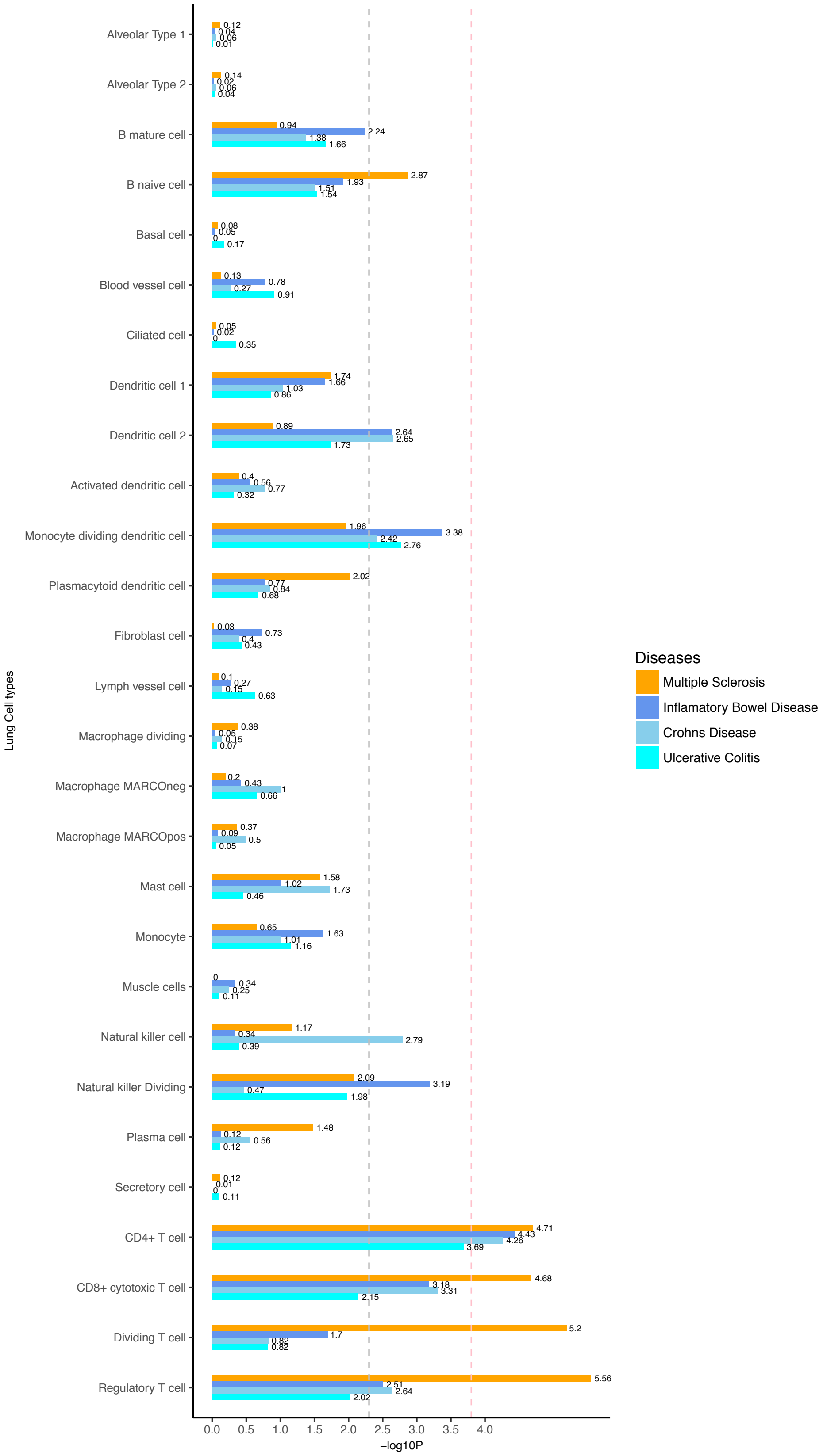

**Figure S13.** Heritability enrichment of 28 lung cell type-specific expressed genes in MS and each of IBD, UC, and CD. Negative log10 p-value of coefficient Z-score are displayed in x axis. The grey and pink dotted line represent the FDR threshold <5% and Bonferroni corrected threshold, respectively.

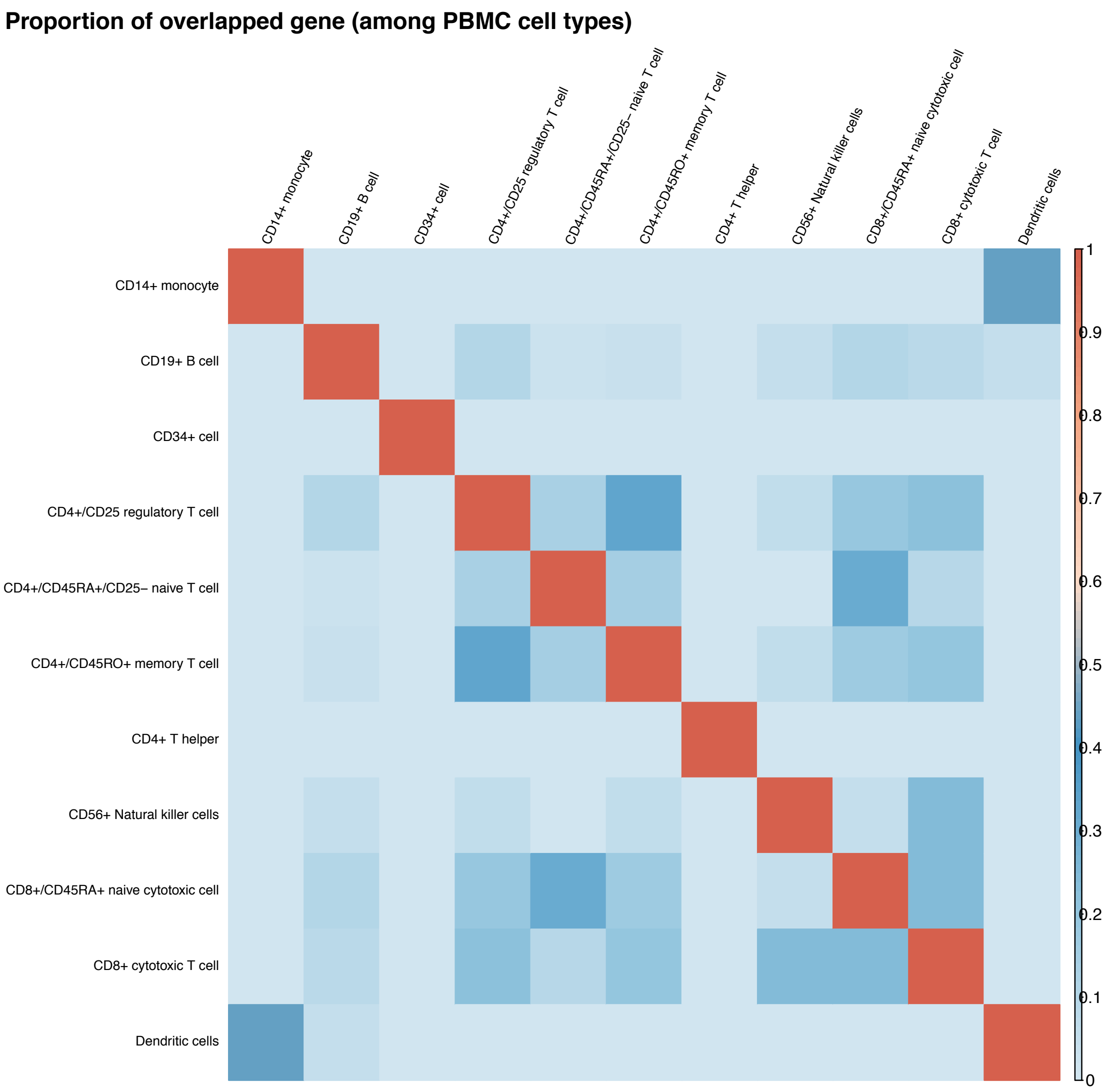

**Figure S14.** Proportion of overlapped top 10% highly expressed genes among 11 peripheral blood mononuclear cells (PBMC) cell types.

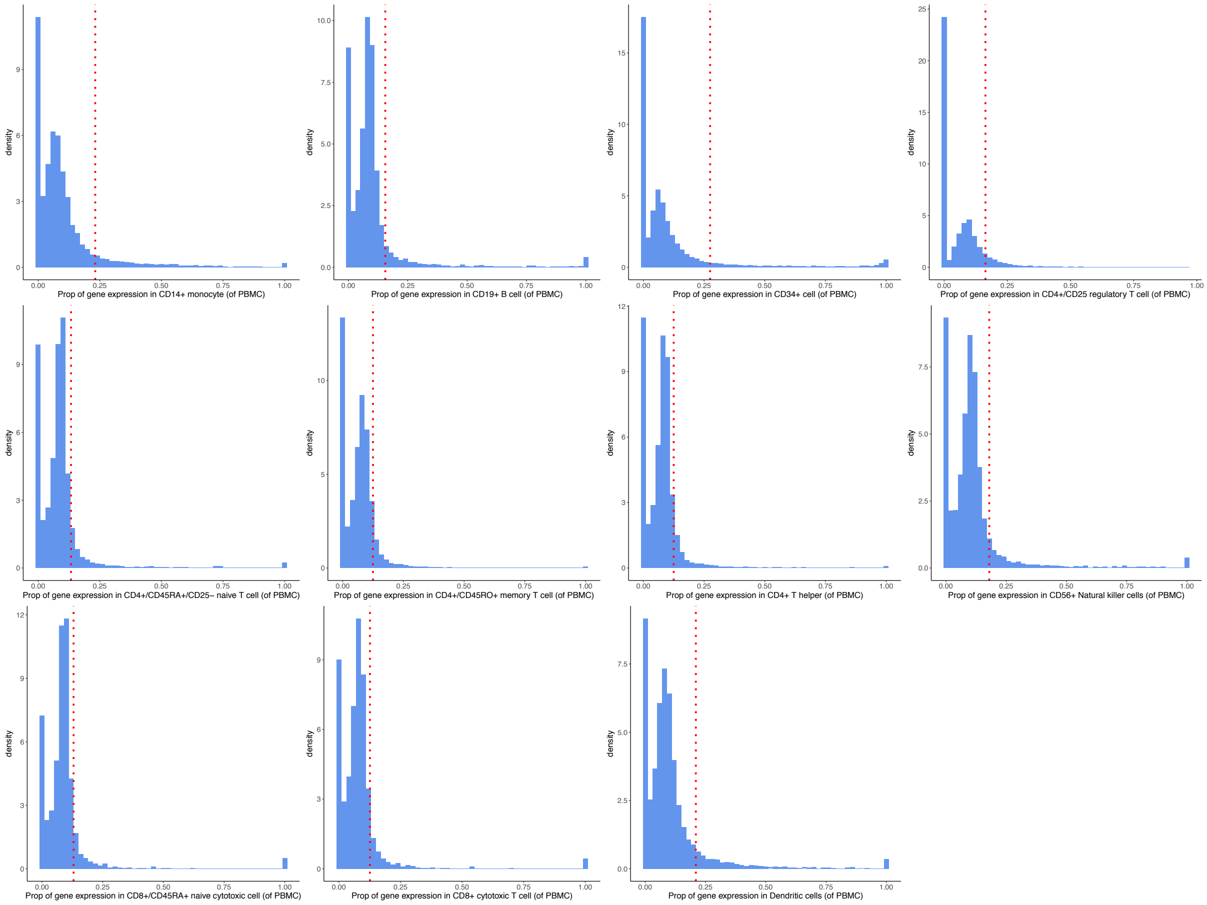

**Figure S15.** Distribution of the proportion of gene expression per cell type in the total gene expressions of all 11 PBMC cell types. The top 10% expressed genes are distributed in the right parts of the red dotted vertical line.

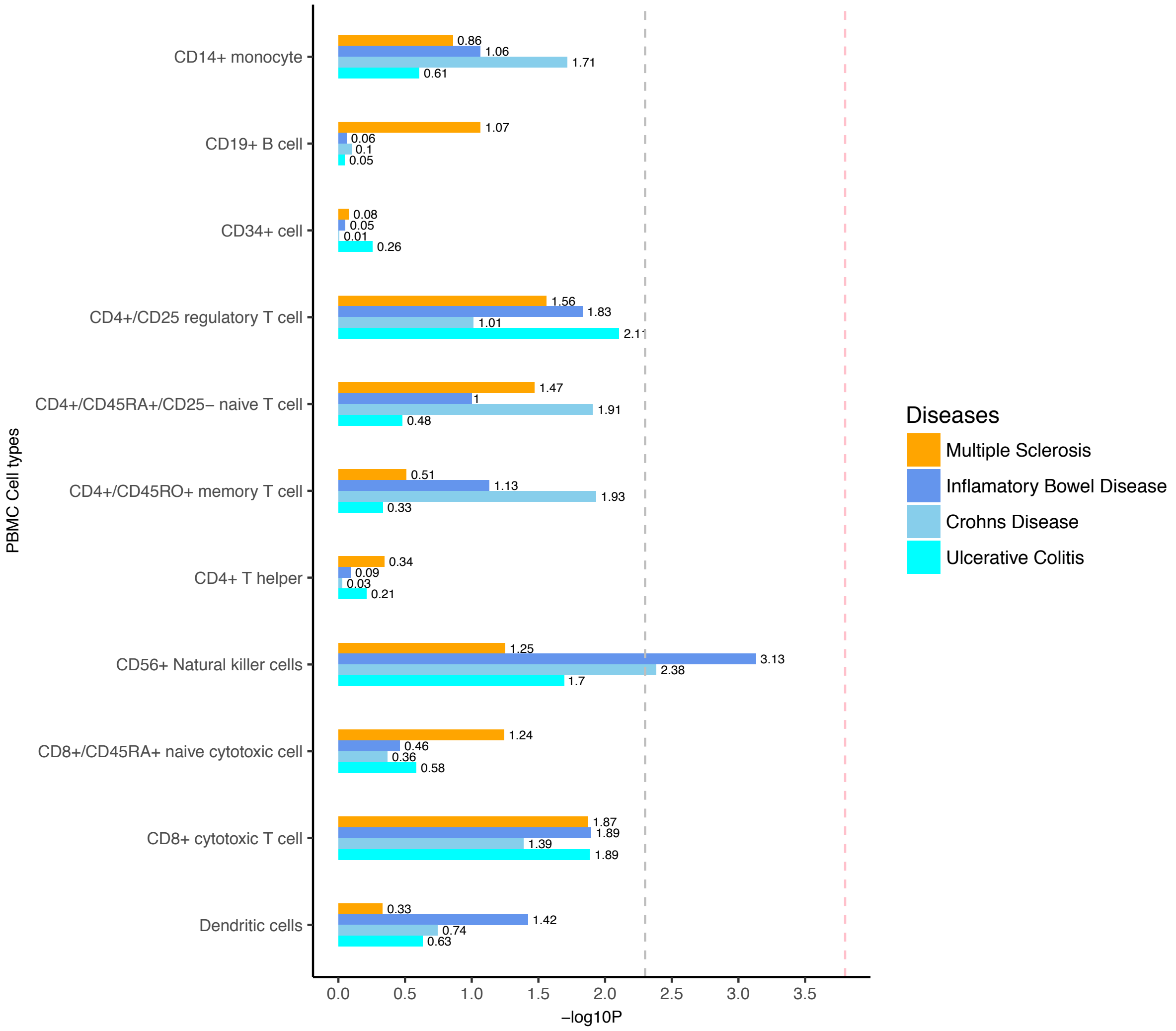

**Figure S16.** Heritability enrichment of 11 PBMC type-specific expressed genes in MS and each of IBD, UC, and CD. Negative log10 p-value of coefficient Z-score are displayed in x axis. The grey and pink dotted line represent the FDR threshold <5% and Bonferroni corrected threshold, respectively.

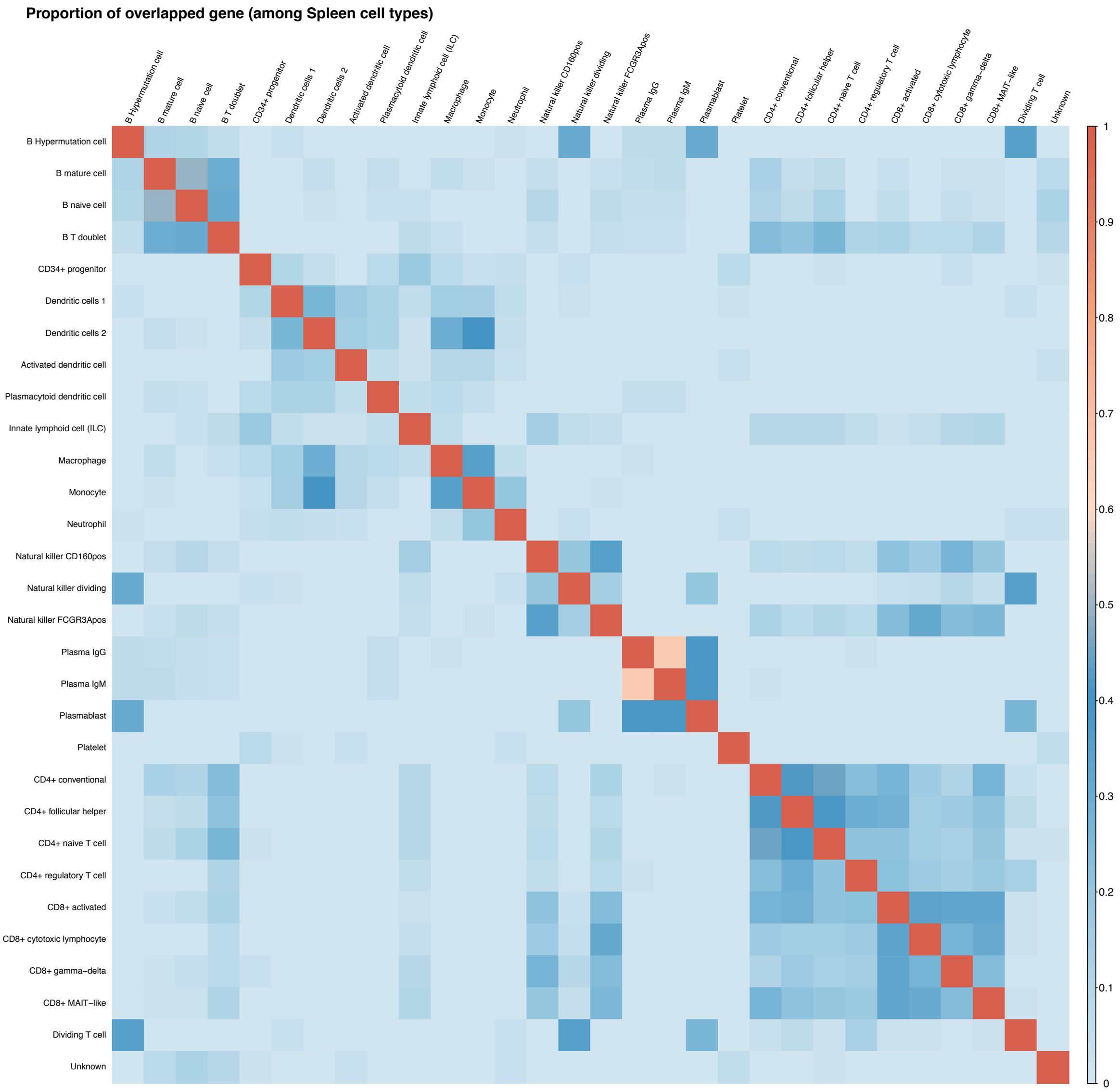

**Figure S17** Proportion of overlapped top 10% highly expressed genes among 30 spleen cell types.

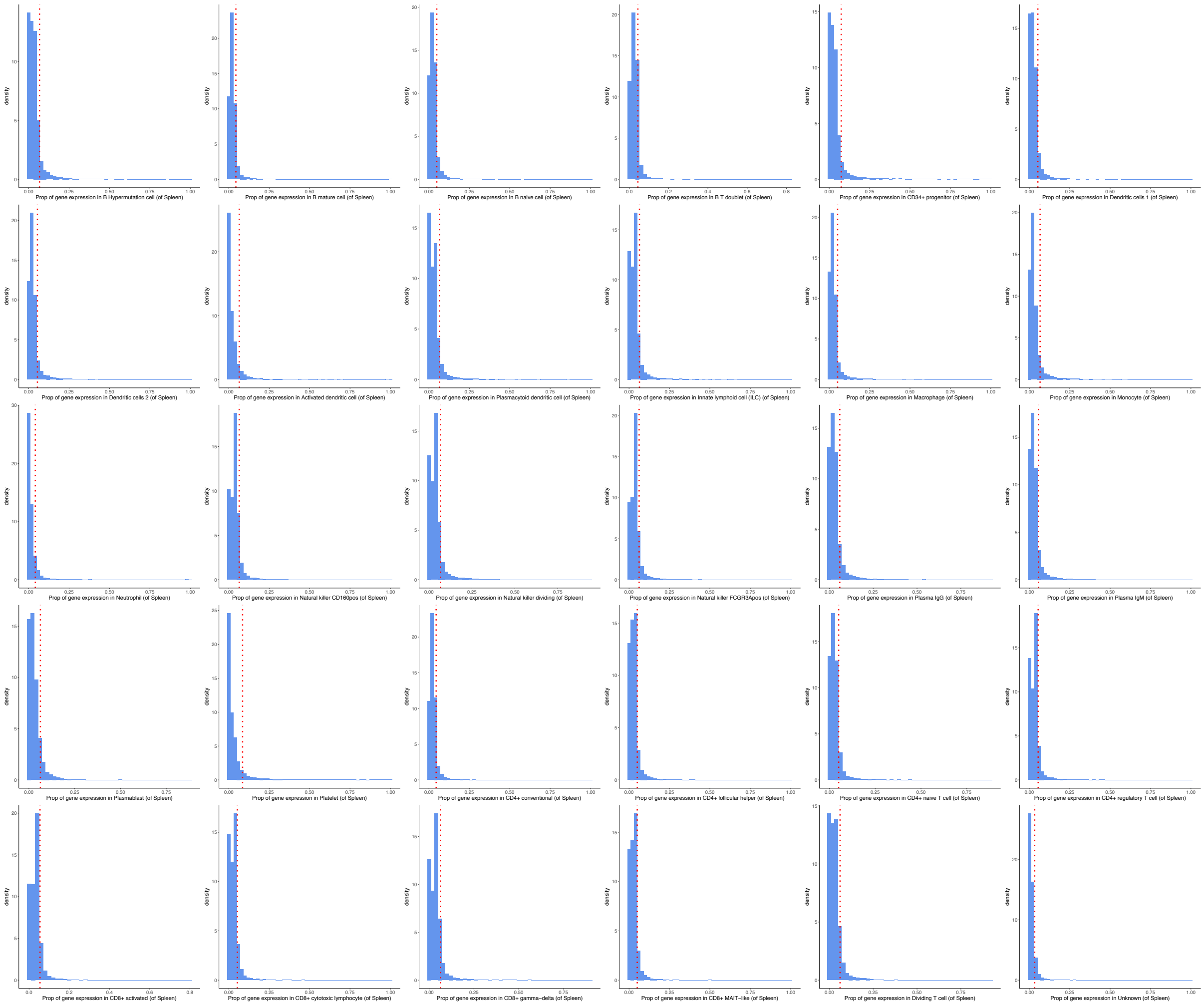

**Figure S18.** Distribution of the proportion of gene expression per cell type in the total gene expressions of all 30 spleen cell types. The top 10% expressed genes are distributed in the right parts of the red dotted vertical line.

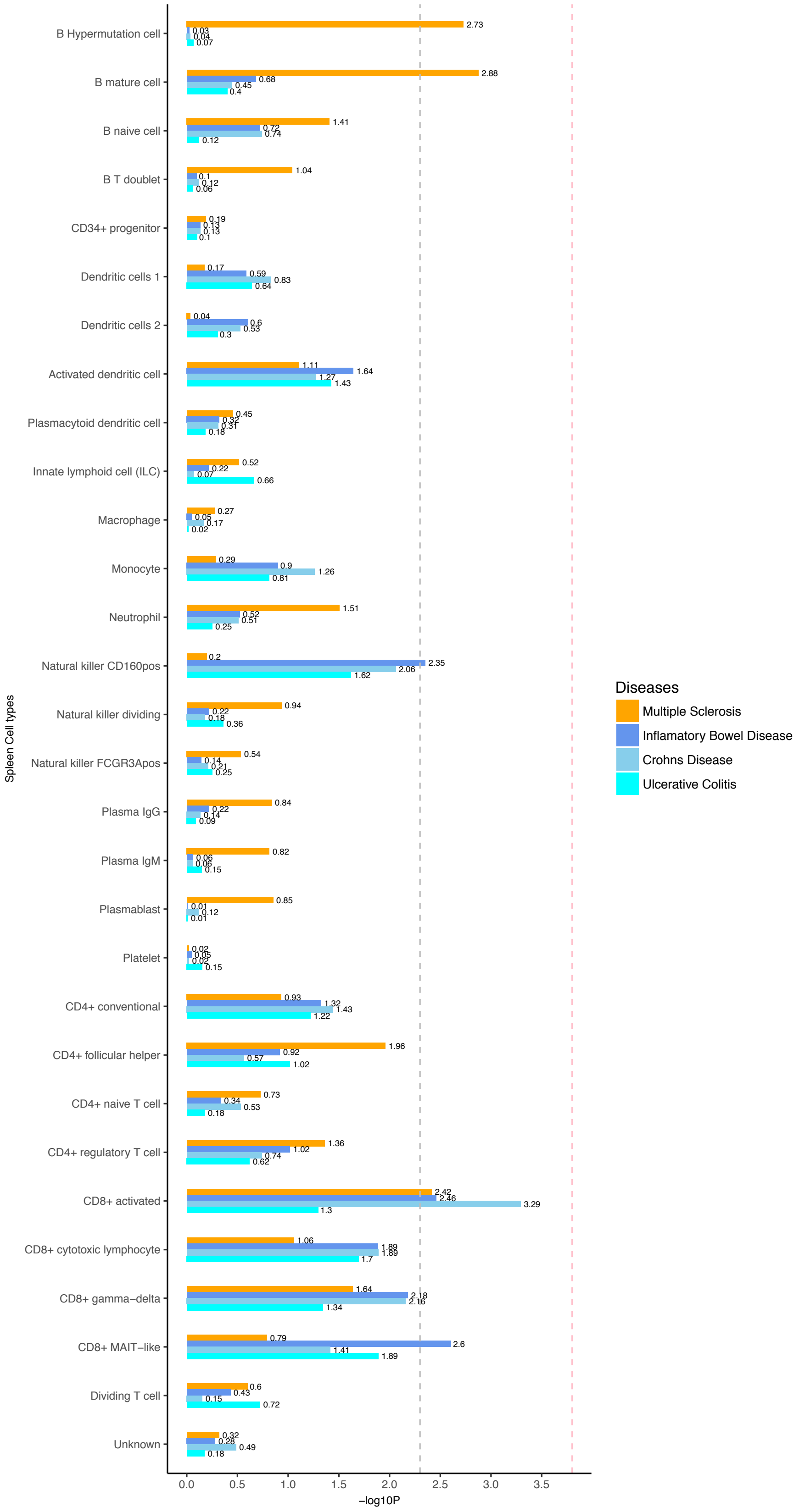

**Figure S19.** Heritability enrichment of 30 spleen cell type-specific expressed genes in MS and each of IBD, UC, and CD. Negative log10 p-value of coefficient Z-score are displayed in x axis. The grey and pink dotted line represent the FDR threshold <5% and Bonferroni corrected threshold, respectively.

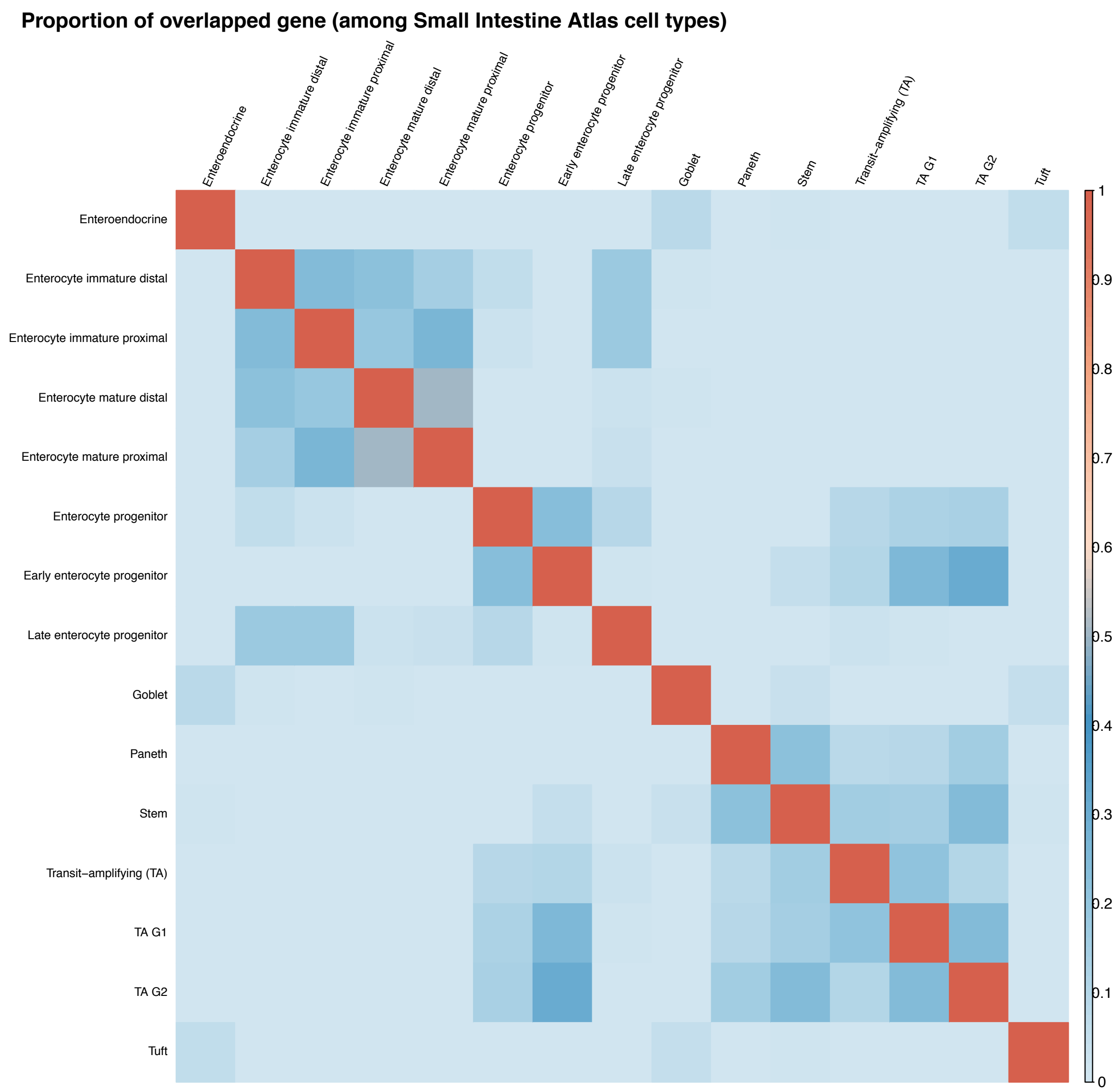

**Figure S20.** Proportion of overlapped top 10% highly expressed genes among 15 small intestine cell types.

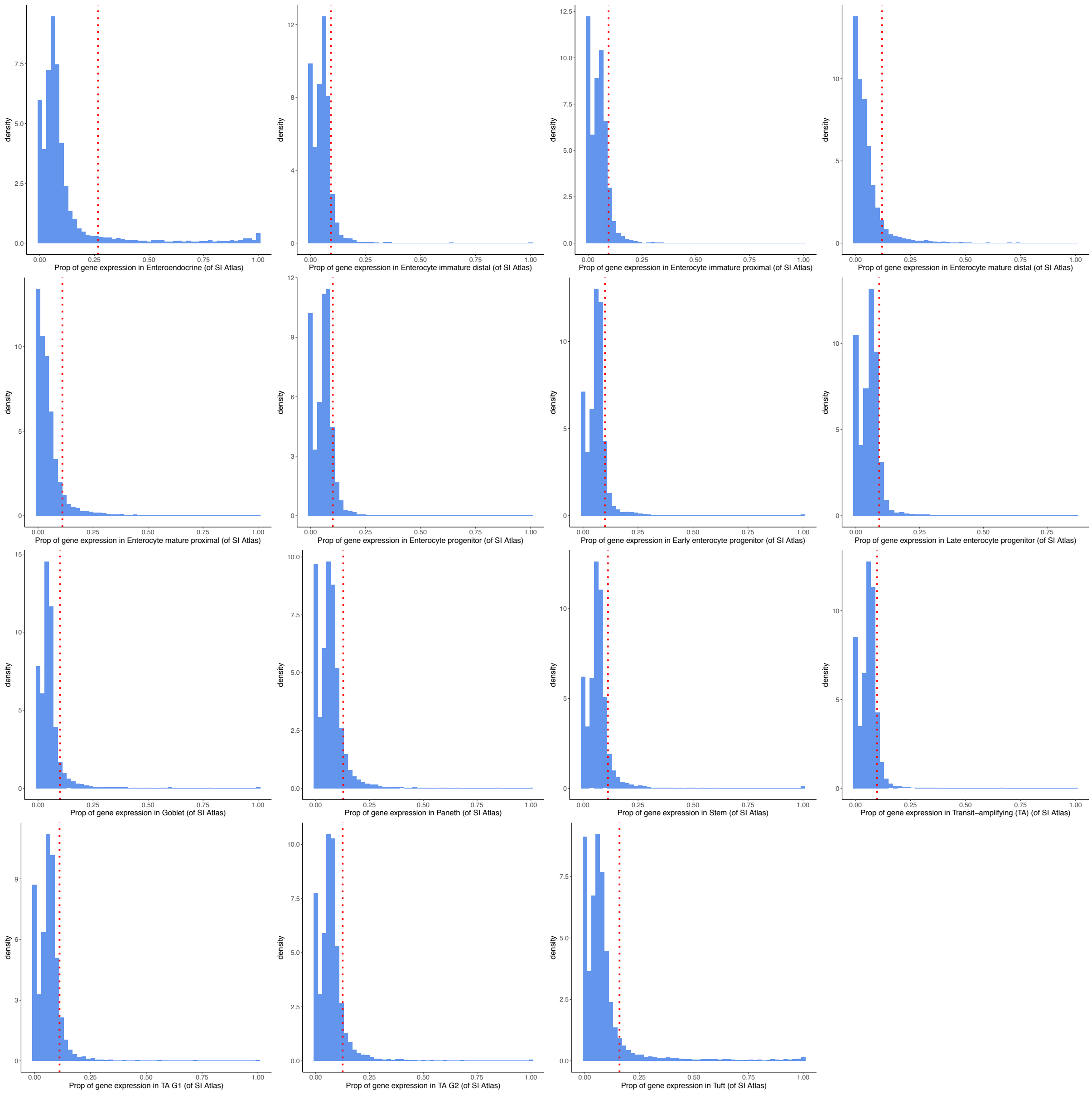

**Figure S21.** Distribution of the proportion of gene expression per cell type in the total gene expressions of all 15 small intestine cell types. The top 10% expressed genes are distributed in the right parts of the red dotted vertical line.

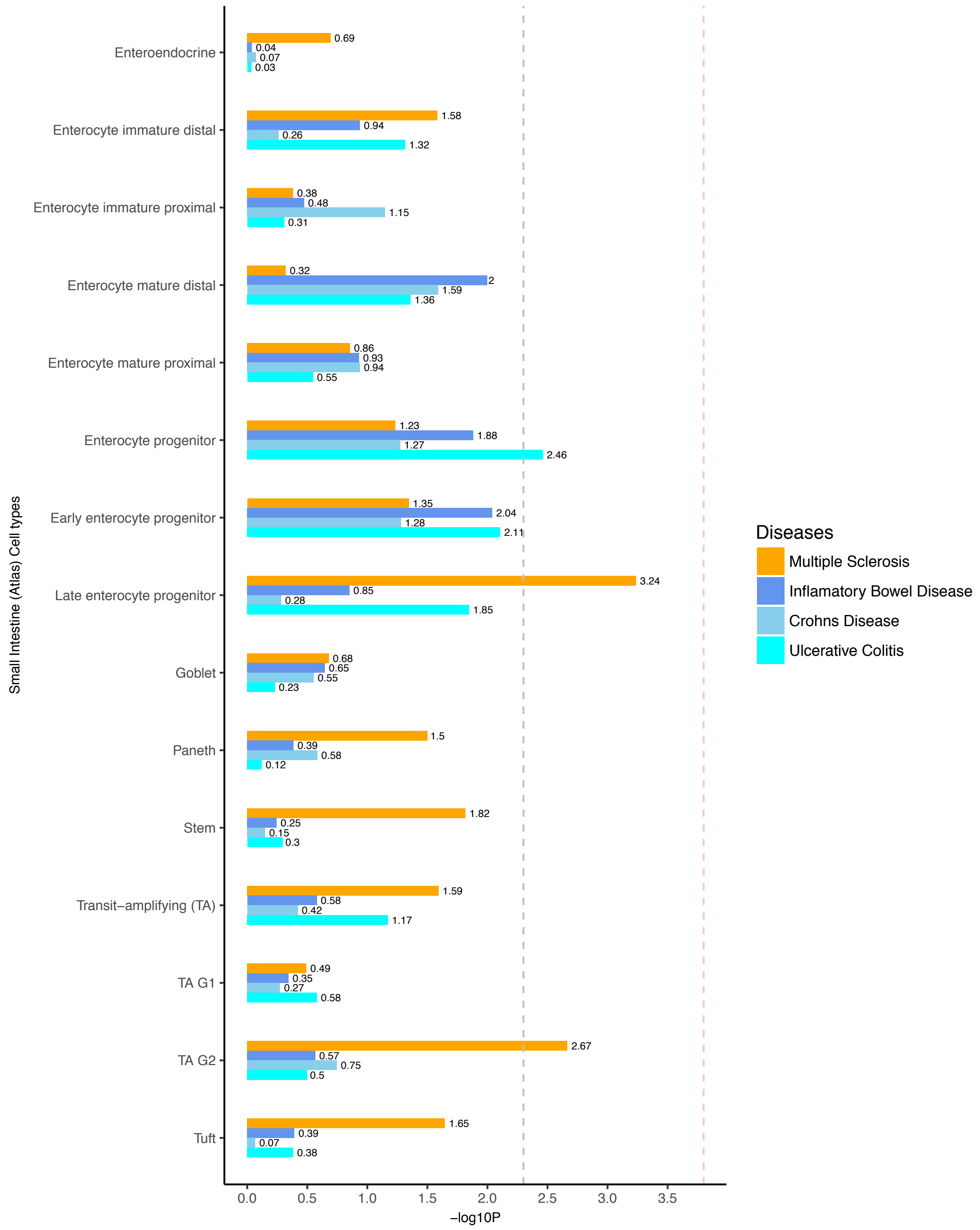

**Figure S22.** Heritability enrichment of 15 small intestine cell type-specific expressed genes in MS and each of IBD, UC, and CD. Negative log10 p-value of coefficient Z-score are displayed in x axis. The grey and pink dotted line represent the FDR threshold <5% and Bonferroni corrected threshold, respectively.

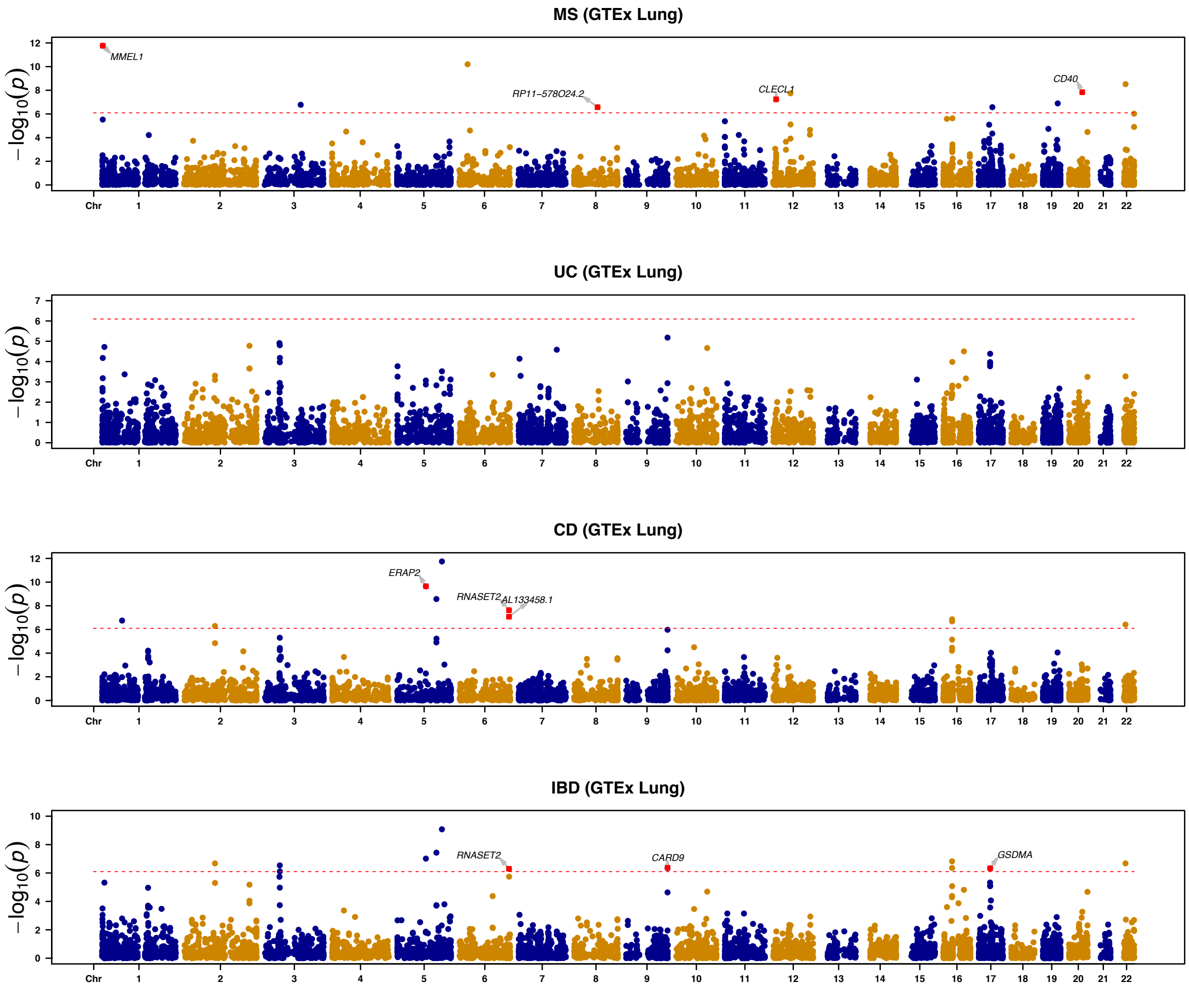

**Figure S23.** Manhattan plots of SMR results for the associations between GTEx (Lung) eQTL summary data and GWAS data of MS and each of IBD, UC and CD. For each plot, y-axis shows the –log10(SMR p-value); the dotted horizontal line indicates the study-wide Bonferroni-corrected SMR threshold (SMR $p < \sim 5.36 \times 10^{-7}$); and the genes in red represent the putative gene expression because of a causal variant with pleiotropic effects, with HEIDI p >0.05 and at least 10 SNPs after HEIDI-outlier test.

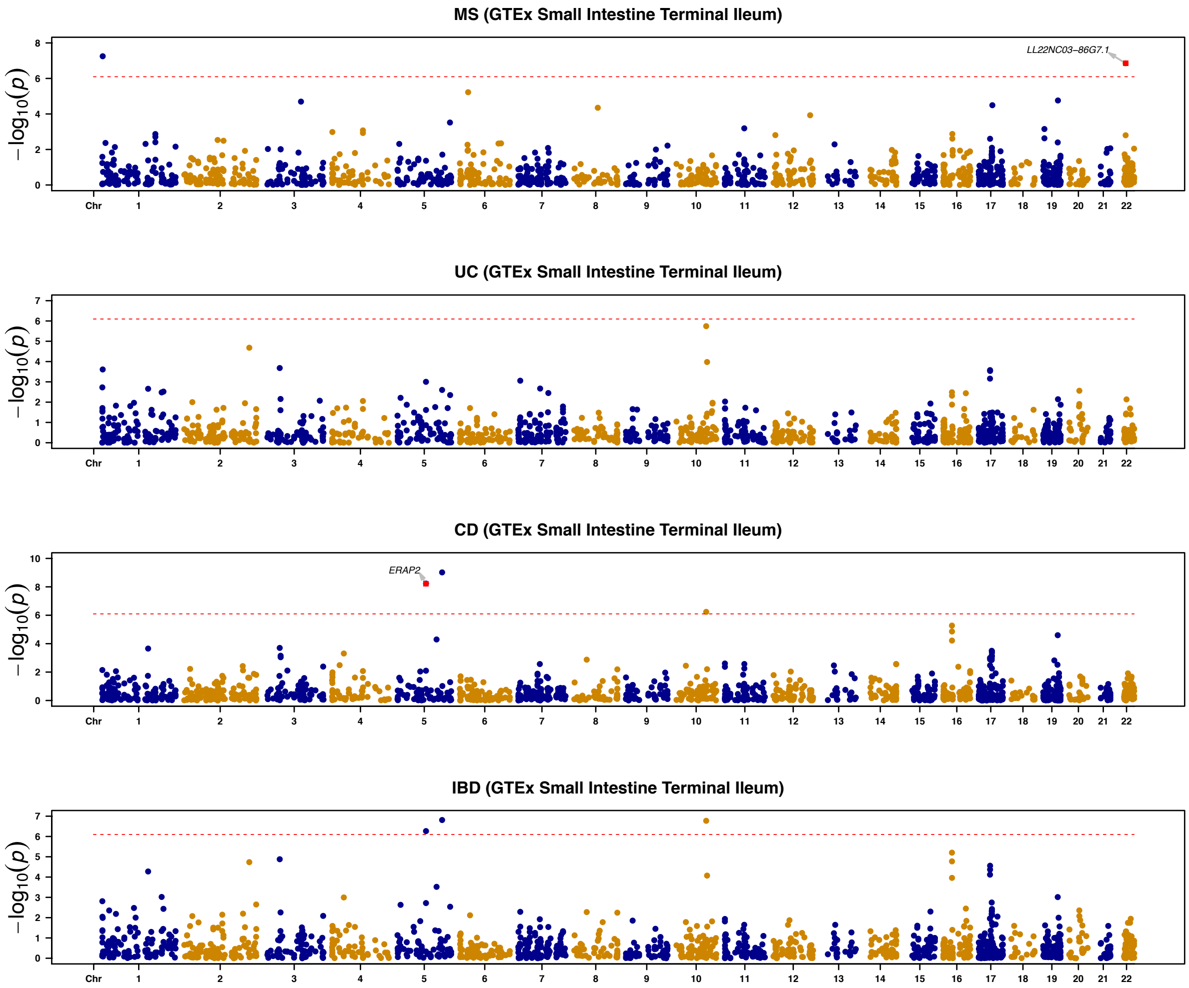

**Figure S24.** Manhattan plots of SMR results for the associations between GTEx (Small Intestine-Terminal Ileum) eQTL summary data and GWAS data of MS and each of IBD, UC and CD. For each plot, y-axis shows the –log10(SMR p-value); the dotted horizontal line indicates the study-wide Bonferroni-corrected SMR threshold (SMR p <~$5.36\times10^{-7}$); and the genes in red represent the putative gene expression because of a causal variant with pleiotropic effects, with HEIDI p >0.05 and at least 10 SNPs after HEIDI-outlier test.

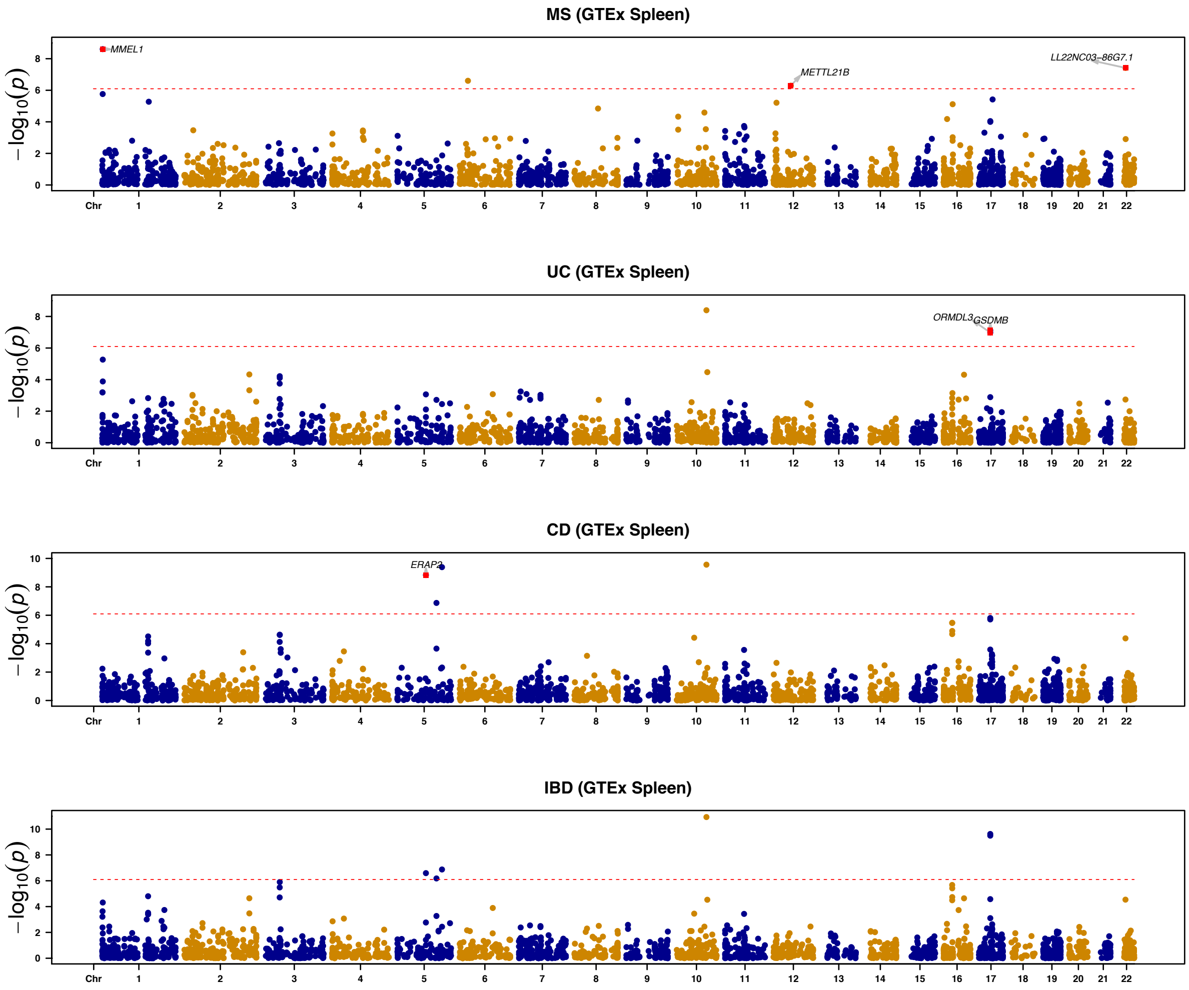

**Figure S25.** Manhattan plots of SMR results for the associations between GTEx (Spleen) eQTL summary data and GWAS data of MS and each of IBD, UC and CD. For each plot, y-axis shows the –log10(SMR p-value); the dotted horizontal line indicates the study-wide Bonferroni-corrected SMR threshold (SMR $p <\sim 5.36\times10^{-7}$); and the genes in red represent the putative gene expression because of a causal variant with pleiotropic effects, with HEIDI p >0.05 and at least 10 SNPs after HEIDI-outlier test.

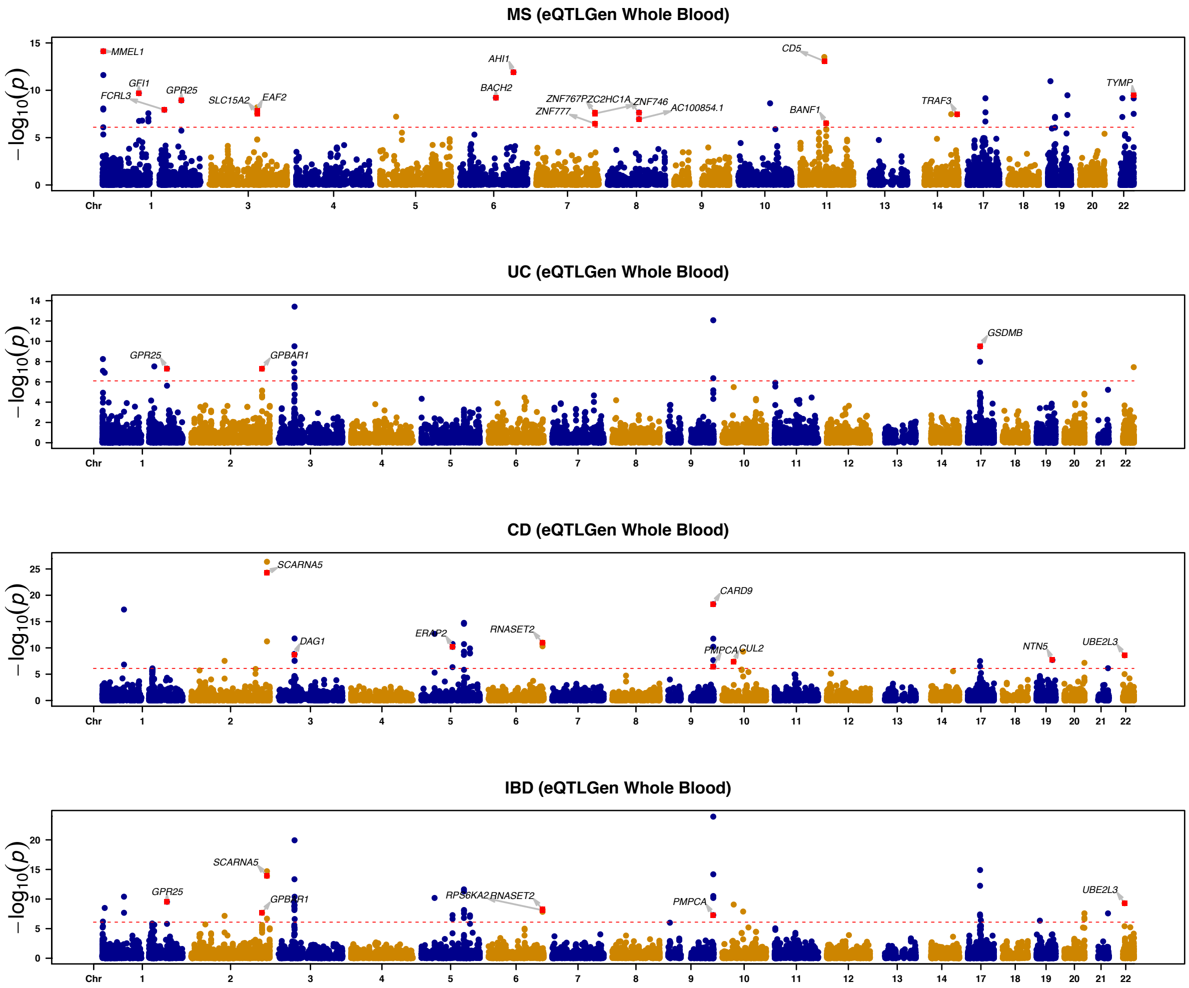

**Figure S26.** Manhattan plots of SMR results for the associations between eQTLGen summary data and GWAS data of MS and each of IBD, UC and CD. For each plot, y-axis shows the –log10(SMR p-value); the dotted horizontal line indicates the study-wide Bonferroni-corrected SMR threshold (SMR p $<\sim 5.36\times10^{-7}$); and the genes in red represent the putative gene expression because of a causal variant with pleiotropic effects, with HEIDI p >0.05 and at least 10 SNPs after HEIDI-outlier test.

#### Reference

1. Finucane, H.K. *et al.* Heritability enrichment of specifically expressed genes identifies disease-relevant tissues and cell types. *Nat Genet* **50**, 621-629 (2018).
2. Bryois, J. *et al.* Genetic identification of cell types underlying brain complex traits yields insights into the etiology of Parkinson's disease. *Nat Genet* **52**, 482-493 (2020).
